# Network predictions sharpen the representation of visual features for categorization

**DOI:** 10.1101/2022.07.01.498431

**Authors:** Yuening Yan, Jiayu Zhan, Robin A.A. Ince, Philippe G. Schyns

## Abstract

Models of visual cognition assume that brain networks predict the contents of a stimulus to facilitate its subsequent categorization. However, the specific network mechanisms of this facilitation remain unclear. Here, we studied them in 11 individual participants cued to the spatial location (left vs. right) and contents (Low vs. High Spatial Frequency, LSF vs. HSF) of an upcoming Gabor stimulus that they categorized. Using concurrent MEG recordings, we reconstructed in each participant the network that communicates the predicted contents and the network that represents these contents from the stimulus for categorization. We show that predictions of LSF vs. HSF propagate top-down from temporal to contra-lateral occipital cortex, with modulatory supervision from frontal cortex. In occipital cortex, predictions sharpen bottom-up stimulus LSF vs. HSF representations, leading to faster categorizations. Our results therefore reveal functional networks that predict visual contents to sharpen their representations from the stimulus to facilitate categorization behavior.

---

Since Helmholtz’s “unconscious inferences”, vision scientists have worked with the hypothesis that what we visually perceive is influenced by the bottom-up sensory input, but also by top-down expectations of what this input might be ^1,2^. Expectations predict upcoming visual information contents ^3–5^, thereby facilitating their disambiguation from the noisy input ^6,7^ to speed up categorization behavior ^8^.

However, specifically where and when brain networks predict visual contents and how these improve subsequent stimulus categorization remain unclear. Advances in neuroimaging showed that top-down predictions modulate the local neural signal in multiple ways, by inducing patterns of stimulus-specific activation with reduced alpha oscillation power, with enhanced gamma-band activity ^9–12^, or with selective changes of cortical layer activity, as shown with high-field fMRI ^13^. However, we do not know the network communications that top-down propagate the predictions of specific visual contents to occipital cortex, nor how these predicted contents in turn change the representation of the stimulus to speed up behavior. One hypothesis suggests that neural predictions of sensory contents sharpen their representations from the stimulus ^7^, while suppressing irrelevant contents ^14^. Conversely, neural predictions could dampen predicted contents ^1,15^, thereby enhancing selectivity to the unpredicted contents ^16,17^. Hence, we still need to tease apart these accounts of the effect of top-down prediction bottom-up stimulus representation.

Here, we addressed three fundamental questions pertaining to the prediction of visual contents and their processing from the stimulus for categorization (illustrated in Figure1):

**Figure 1.**
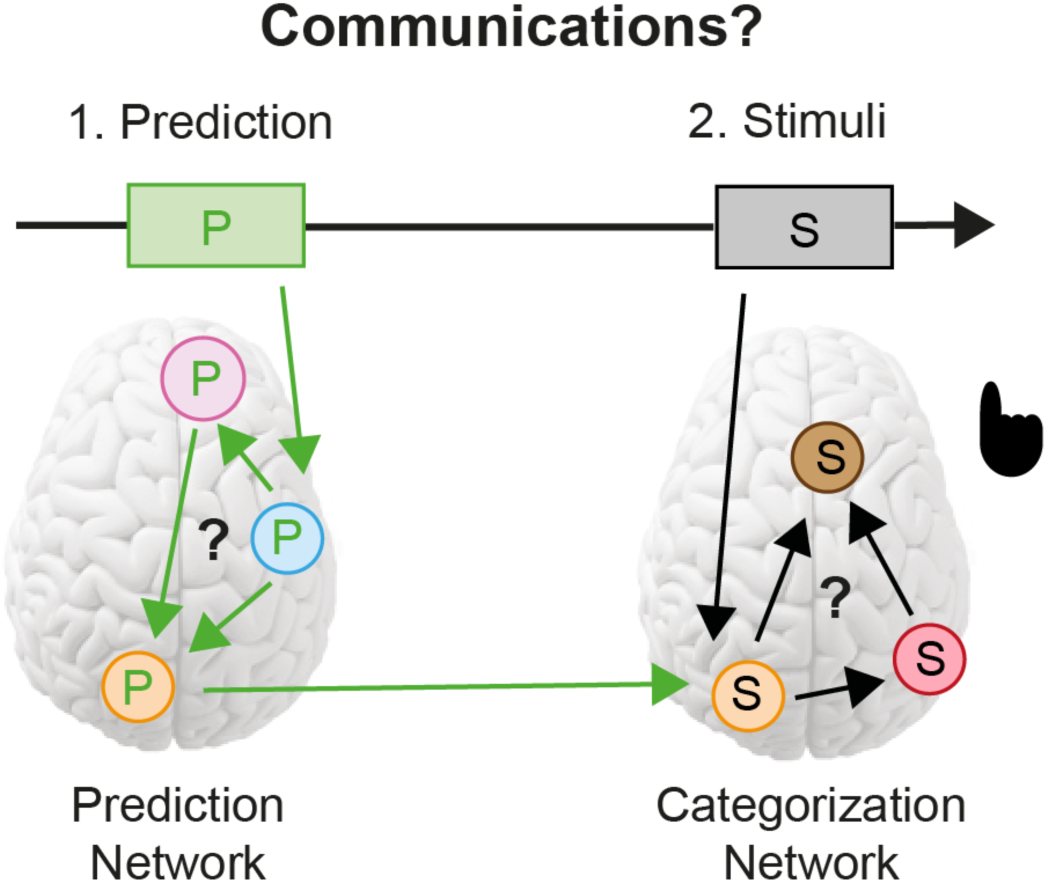
Three key questions. **1) Prediction Network:** Where, when and how does it communicate a predictive cue (here P, an auditory cue that predicts a visual content). **2) Categorization network:** When, where and how does it process the stimulus contents? **3) Influence of prediction on categorization:** How does prediction influence stimulus processing leading to faster behavior?

1. When, where, and how does a prediction of stimulus contents dynamically propagate through a Prediction Network of brain regions?
2. When, where, and how do the contents of a presented stimulus dynamically propagate in a Categorization Network for behavior?
3. How does content predictions in (1) change stimulus content representations in (2) to speed up behavior?

## Results

### Cueing speeds up behavior

Our three-stage cueing design is depicted in Figure 2A and 2B. On each trial, a spatial cue at Stage 1 (green dot briefly displayed left vs. right of a central fixation cross (cf. Posner cueing ^18^) predicted the visual hemifield location (left vs. right) of an upcoming Gabor patch (henceforth, Gabor, see *Methods, Stimuli*) with 100% validity, followed by a 1-1.5s blank screen. At Stage 2, on informative trials an auditory cue (a 250ms sweeping tone at 196 Hz vs. 2217 Hz) predicted the Spatial Frequency content (SF, Low vs. High) of the upcoming Gabor stimulus with 90% validity (henceforth, refer as the “predicted” trials), followed by another 1-1.5s blank interval. On uninformative, “non-predicted” trials (33% of Stage 2 trials), a 622 Hz neutral auditory cue had no association with LSF or HSF. Finally, at Stage 3, the actual Gabor stimulus appeared in the participant’s left vs. right visual hemifield for 100ms. Each participant (N = 11, see *Methods, Participants*) categorized the Gabor SF as quickly and accurately as they possibly could without feedback (i.e. 3-AFC, with responses “LSF” vs. “HSF” vs. “don’t know”, see also *Methods, Procedure*).

**Figure 2.**
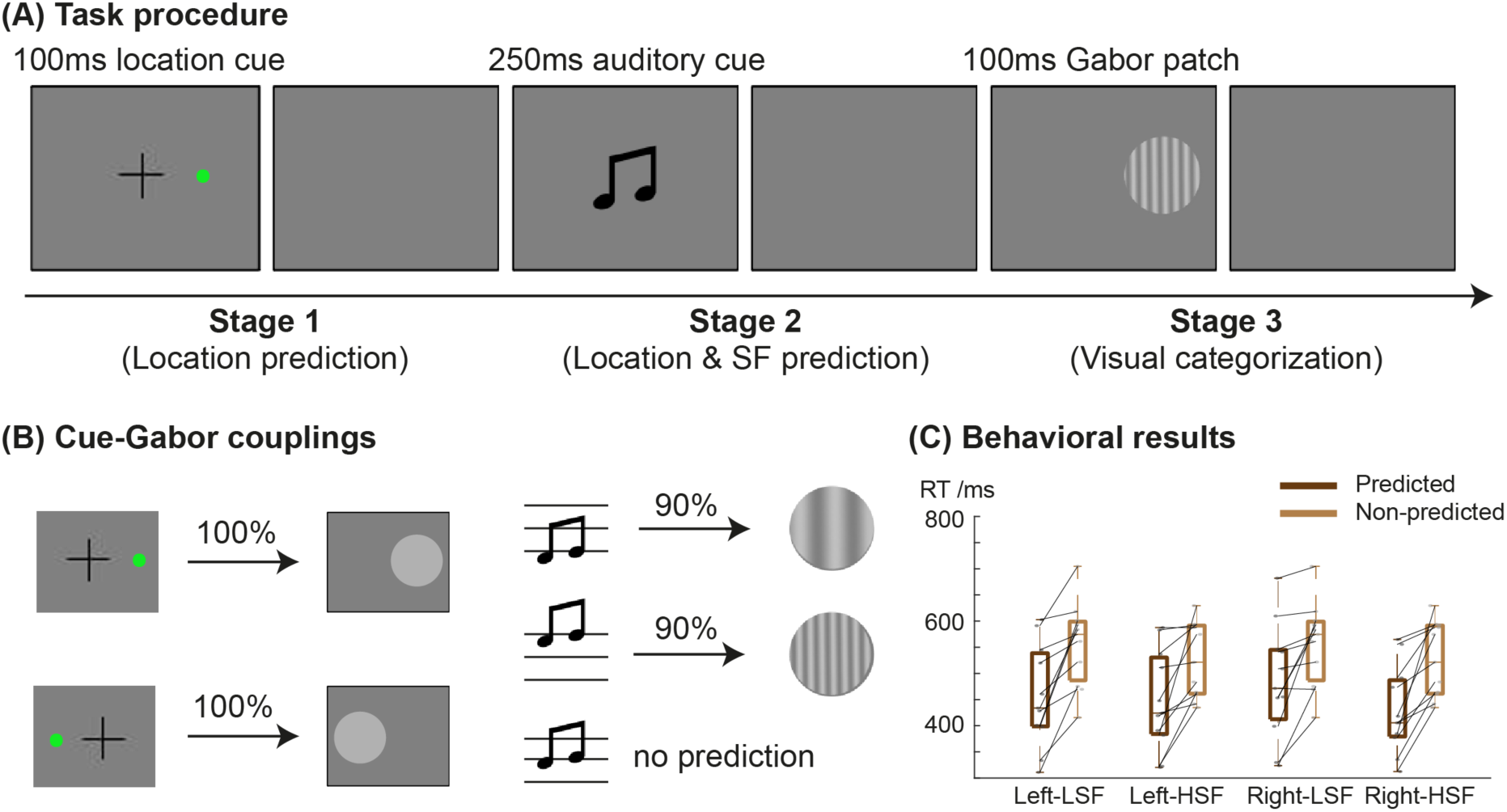
Experimental design and behavioral results. **(A) Task procedure.** Each trial started with a 500ms fixation. At Stage 1, a 100ms green dot (i.e. location cue) predicted the location of the upcoming Gabor patch, followed by 1000-1500ms blank screen. At Stage 2, a 250ms sweeping sound (i.e., SF cue) predicted the LSF vs. HSF content of the upcoming Gabor, followed by a 1-1.5s blank screen with jitter. At Stage 3, the Gabor stimulus was presented for 100ms. Participants categorized its LSF vs. HSF, followed by a 750-1250ms inter-trial interval (ITI). **(B) Cue-Gabor couplings**. At Stage 1, the left vs. right location cue predicted the left vs. right location of the upcoming Gabor with 100% validity. At Stage 2, the 196 Hz vs. 2217 Hz auditory cues predicted the Gabor LSF vs. HSF contents with 90% validity; a 622 Hz neutral auditory cue served as non-prediction control on 33% of the trials, (i.e. with .5 probability of LSF vs. HSF). **(C) Behavioral results**. Boxplots show that prediction (i.e., valid informative cueing, dark brown) sped up median LSF vs. HSF Gabor categorization RTs in left and right-presented trials, compared with non-prediction (i.e., neutral cueing, light brown). Black dots (vs. light grey dots) indicate the per-participant median categorization RTs in predicted (vs. non-predicted) trials, linked to show directional RT differences replicated in each individual participant.

As expected, valid predictive cues (i.e., the 90% valid trials with informative cueing) improved categorization accuracy compared to non-predicted (neutral cue) trials, on average by 2.58% (96.9% vs. 94.3%), *F(1,86)*=22.5, *p*=0.0008, and sped up Reaction Times (RTs), on average by 87.7ms (454.4ms vs. 542.1ms), *F(1, 86)*=20.8, *p*=0.001. Significant RT improvements applied to each Gabor location×SF presentation condition (see Figure 2C, Supplemental Table S1 and *Methods, Cueing improves behavior*) and to each individual participant (Bayesian population prevalence ^19,20^ 11/11 = 1 [0.77 1] (Maximum A Posteriori (MAP) [95%, Highest Posterior Density Interval (HPDI)], see Supplemental Table S2).

Behavioral facilitation therefore validates our cueing design and warrants the study of the network mechanisms whereby Stage 2 prediction influence Stage 3 categorization. To do so, we first reconstruct the Prediction and the Categorization networks in each participant.

### Reconstructing the Stage 2 Prediction Network

We reconstructed the concurrently measured participant’s MEG activity on 12,773 sources, see *Methods, MEG Data Acquisition and Pre-processing*. To identify the brain regions that propagate the prediction prior to stimulus onset (cf. Figure 1), we first computed how strongly each MEG source dynamically represents the prediction at Stage 2 (when participants can integrate the location prediction from Stage 1), following cue onset—i.e. as the Mutual Information (MI) between LSF vs. HSF auditory cue and Stage 2 MEG source activity, on each source and time point, MI(LSF vs. HSF auditory cue; Stage 2 MEGt). Figure 3 presents these prediction representation data at source level, color-coded by source representation onset time at Stage 2.

**Figure 3.**
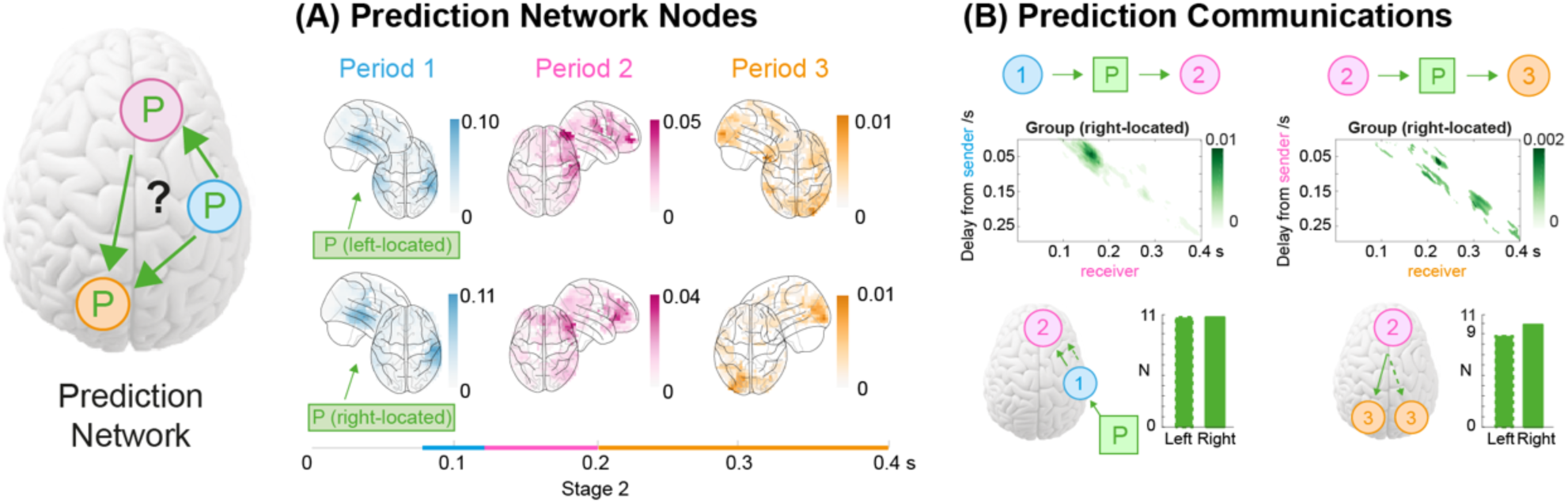
Stage 2 Prediction Network. **(A) Network nodes (see iconic brain).** To identify the nodes, in each participant, separately for left-vs. right-located trials, we computed the prediction representation (as MI(LSF vs. HSF cue; MEG_*t*_), Y-axis), between 0 and 0.4s post auditory cue (X-axis), on each of 4,413 sources (see *Methods, Prediction representations*). We then localized sources with peak MI in each period—i.e. [90-120ms], [120-200ms], [>200ms], see *Methods, Prediction Network, Prediction periods clustering*. Glass brain shows the cross-participant mean of peak MI for these sources, revealing the temporal sequence of prediction from temporal (cyan) to frontal (magenta) to occipital (orange, contra-laterally to the left vs. right predicted stimulus location). **(B) Prediction communications**. For each participant, we used the source with peak prediction representation MI in each space x time region as the three functional nodes of the Prediction Network. We used DFI ^21^ to compute Stage 2 communications of the prediction across these network nodes— i.e. 1(temporal)->2(frontal) and 2(frontal)->3(occipital). Plots show the strength of communications averaged across participants, in the time course of the receiving node (X-axis), as communication delays from the sending node (Y-axis). For example, we can see that temporal node 1 sends the predictive cue P to frontal node 2, with a 50-100 ms delay (as illustrated in the iconic brain below). Individual participant replications of 1->2, left-located trials (dashed) 11/11, right-located trials 11/11, Bayesian population prevalence = 1 [0.77 1] MAP [95% HPDI]; 2->3, left-located trials, 9/11, Bayesian population prevalence = 0.81 [0.53 0.96], MAP [95% HPDI], right-located trials,10/11, Bayesian population prevalence = 0.91 [0.64 0.99], MAP [95% HPDI].

Figure 3 shows that the prediction propagates over three distinct periods (revealed by a clustering analysis, see *Methods, Prediction periods clustering* and Supplemental Figure S1). Figure 3A illustrates each period, by locating in each participant (N = 11, see Supplemental Figure S2 for individual results) the source that maximally represents the prediction in each time window (i.e. peak MI), separately for left-vs. right-located trials (see *Methods, Prediction network nodes*). Prediction representation starts with an early peak in the temporal lobe (i.e., auditory cortex, Period 1), moving to frontal cortex ([120-200ms], magenta, Period 2), and then to the occipital cortex contra-lateral to the predicted location ([>200ms], orange, Period 3). Additionally, prediction propagation is distinct from that of the bottom-up auditory input, as tested on the localiser data prior to the cueing task (see Supplemental Figure S3 and *Methods, Auditory localizer* and *Auditory localizer representation*).

Representation dynamics suggest a functional Prediction Network that communicates the predictive cue. To reconstruct this network, we measured the communications of Prediction P across the just identified regions (i.e. temporal, frontal and occipital)—computed with Directed Feature Information ^21^, as DFI_P_(regionX→regionY), FWER-corrected, *p*<0.05 (see *Methods, Prediction network reconstruction* and Supplemental Figure S4). Figure 3B shows these communications for right-located trials, whose sequence starts in temporal lobe, propagating to frontal cortex (i.e. PFC, in the dIPFC region) and then on to occipital cortex. Prevalence bars in Figure 3B indicate high individual participant replications of the Prediction Network communications—in at least 9/11 participants, Bayesian population prevalence = 0.81 [0.53 0.96], MAP [95% HPDI], see Supplemental Figure S5 for all individual data including left-located trials.

Next, we test the role of frontal cortex in communicating the predictive cue from temporal to occipital cortex (see *Methods, Prediction network modulation*). Figure 4 contrasts a direct communication from temporal to occipital cortex, without frontal mediation (Figure 4A), to a communication mediated by frontal cortex (Figure 4B). We show in Figure 4 that communications of the predictive cue from temporal to occipital regions (i.e. 1->3) is conditional on frontal source 2 activity, which therefore mediates the communication—independently replicated for left- and right-located trials in at least 10/11 participants, individual results in Supplemental Figure S6, Bayesian population prevalence = 0.91 [0.64 0.99] (MAP [95% HPDI]). We conclude that PFC (dIPFC) mediates network communications of the prediction from temporal to occipital cortex.

**Figure 4.**
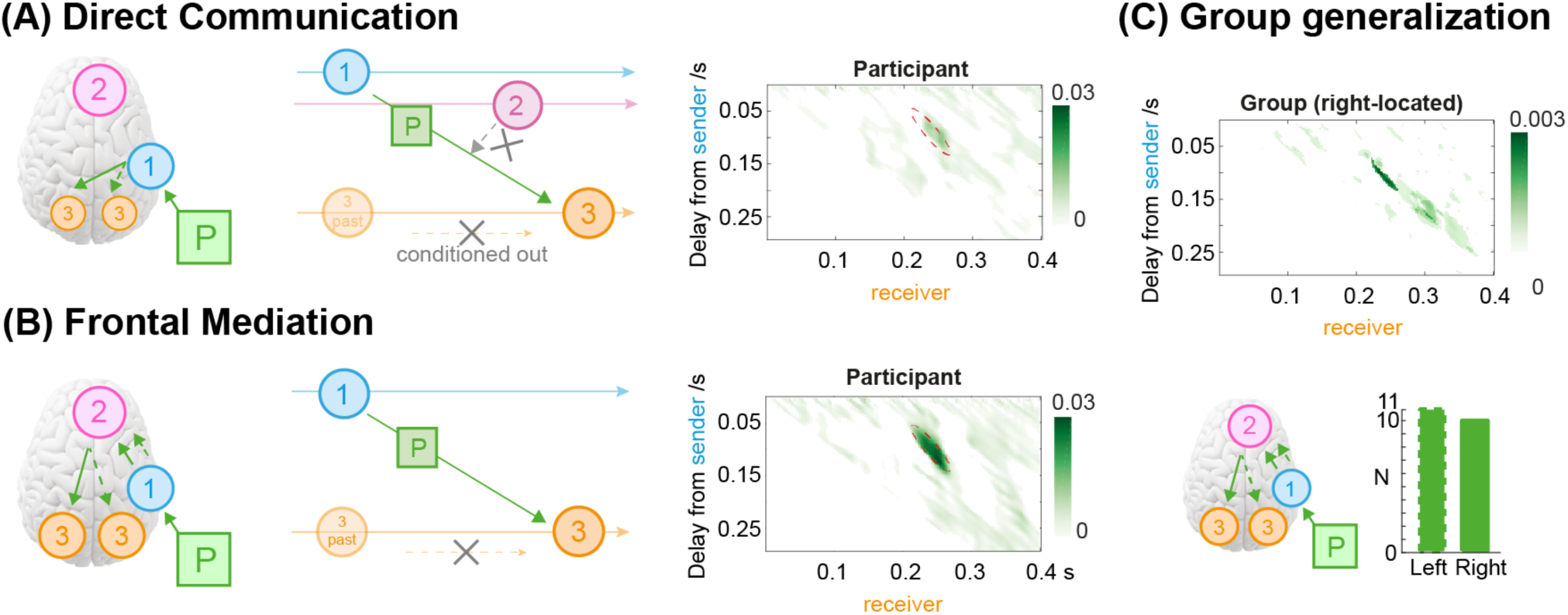
Mediation of prediction communication in frontal cortex. **(A) Direct communication** of the prediction from temporal (source 1) to occipital cortex (source 3). We condition out (i.e. remove) the modulatory role of frontal cortex (source 2) to communicate the prediction between source 1 and source 3. This is expressed in the time course of the receiving source 3 time course (X-axis), as communication delays from the sending source 1 (Y-axis). **(B) Mediation of frontal cortex** (source 2) to communicate the prediction from temporal (source 1) to occipital cortex (source 3). Illustrated in a typical participant showing a 100-150 ms delay between source 1 sender and source 3 receiver. The significant difference between (A) Direct and (B) Mediated communications is indicated as the red dashed line in the participant plot (FWER corrected, *p*<0.05, one-tail). **(C) Group generalization**. The plot shows the cross-participant mean of the significant difference between (A) Direct and (B) Mediated communication for right-located trials. We replicated the effect in 11/11 participants for left-located trials (FWER corrected, *p*<0.05, one-tail), Bayesian population prevalence = 1 [0.77 1] MAP [95% HPDI], and in 10/11 participants for right-located trials, Bayesian population prevalence = 0.91 [0.64 0.99], MAP [95% HPDI]. Frontal cortex therefore mediates network communications of the prediction from temporal to occipital cortex.

### Reconstructing the Stage 3 Categorization Network

We have reconstructed the Stage 2 Prediction network. Here, we show that these predicted contents then modulate the representation of the stimulus, when it is shown at Stage 3, in a functional network that categorizes the stimulus.

We reconstructed a Categorization Network as we reconstructed the Prediction Network. In a first step, we used MI to compute the Stage 3 dynamic representation of the Gabor stimulus—i.e. as MI(Gabor SF; Stage 3 MEG_t_), on each source and time point. A clustering analysis revealed again three sequential space x time periods of stimulus representation (see *Methods, Categorization periods clustering* and Supplemental Figure S1). Figure 5A presents these periods at source level, again color-coded by ranked peak LSF vs. HSF representation (MI) time. Stimulus representation starts with an early occipital-ventral LSF vs. HSF representation peak contra-lateral to the presented Gabor stimulus ([150-250ms], orange, Period 4), followed by a parietal lobe peak ([250-350ms], red, Period 5) and a premotor-frontal cortex peak ([> 350ms], brown, Period 6), independently for left and right located (see *Methods, Categorization Network, Categorization periods clustering* and Supplemental Figure S7 for all individual results).

**Figure 5.**
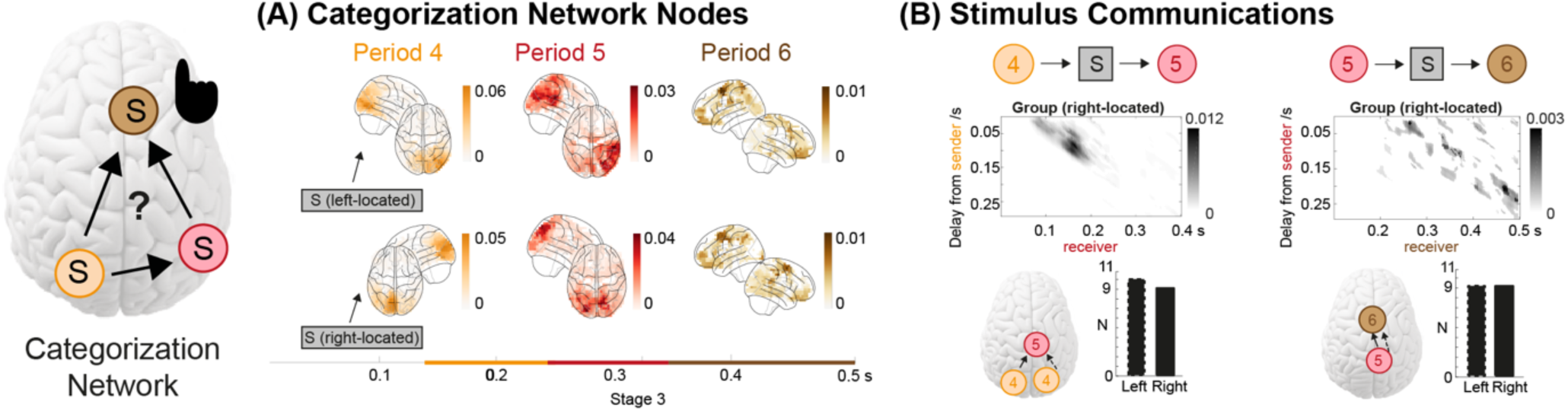
Stage 3 Categorization Network. **(A) Network nodes (see iconic brain)**. To identify the categorization network nodes, in each participant, separately for left-vs. right-located trials, we computed the stimulus representation (as MI(LSF vs. HSF Gabor; MEG_*t*_), Y-axis) between 0 and 0.5s post stimulus, X-axis, on each of 4413 sources. We then localize sources that peak in each period— i.e. [150-250 ms], [250-350 ms], [>350 ms]. Glass brain shows the cross-participant mean peak MI for these sources, revealing the temporal sequence LSF vs. HSF representation starting in contra-lateral occipital-ventral cortex (orange), then parietal regions (red), and finally premotor and frontal cortex (brown), independently for left-located and right-located trials. **(B) LSF vs. HSF communications**. For each participant, we used these three peak MI sources for each space x time region as the three functional nodes of the Categorization Network. We used DFI ^21^ to compute the Stage 3 communications of the stimulus contents across these network nodes—i.e. 4(occipital-ventral)->5(parietal)->6(pre-motor). Plots show the strength of stimulus contents communications averaged across participants, in the time course of the receiving node (X-axis), as communication delays from the sending node (Y-axis). For example, we can see that occipital node 4 sends stimulus S contents to node parietal node 5, with a 75-150 ms delay (as illustrated in the iconic brain plots below). Individual participant replications of 4->5, left-located trials (dashed) 10/11, Bayesian population prevalence = 0.91 [0.64 0.99], MAP [95% HPDI], right-located trials 9/11, Bayesian population prevalence = 0.81 [0.53 0.96], MAP [95% HPDI], 5->6, left-(dashed) and right-located trials 9/11 participants, Bayesian population prevalence = 0.81 [0.53 0.96], MAP [95% HPDI].

Then, with DFI we reconstructed the Categorization Network that communicates the stimulus contents leading to behavior (see *Methods, Categorization network* reconstruction). Figure 5B shows that Gabor LSF vs. HSF communications proceed sequentially from contra-lateral occipital-ventral cortex to parietal and then on to premotor cortex at the average group level. We replicated this Categorization network independently, for left-(dashed lines, Figure 5B) and right-located trials, in ≥ 9/11 participants—i.e. Bayesian population prevalence = 0.81 [0.53 0.96], MAP [95% HPDI], individual results in Supplemental Figure S8.

### Influences of Stage 2 prediction on Stage 3 stimulus categorization

We now turn to how Stage 2 predictions of the upcoming LSF vs. HSF stimulus contents interact with their Stage 3 representations and categorizations categorizations when the Gabor stimulus is shown. To set the stage, Supplemental Figure S9 shows that occipital cortex LSF vs. HSF predictions at Stage 2 preview Stage 3 stimulus representations (see also *Methods, Prediction overlaps with stimulus representation* ^22,23^). It also shows that pre-motor cortex activity on the Categorization Network relates to behavioral RTs (see *Methods, Prediction modulates source activity and RT*). We can now test whether Stage 2 predictions sharpen Stage 3 stimulus representation throughout the categorization network for categorization behavior.

Remember that Stage 2 comprises both informative and neutral trials (cf. Figure 2B, “no prediction”). We can therefore compare in each participant how prediction vs. no prediction changes the source-level representation of the same physical LSF vs. HSF stimulus—computed as MI (LSF vs. HSF; Stage 3 MEG_t_). Figure 6 shows that Stage 2 prediction sharpens Stage 3 discrimination of the predicted LSF vs. HSF contents in contra-lateral occipital (orange), parietal (red), and premotor (brown) regions of the Categorization Network, over the time course of the categorization process (FWER, *p*<0.05, two-tailed, see *Methods, Prediction sharpens stimulus representation*).

**Figure 6.**
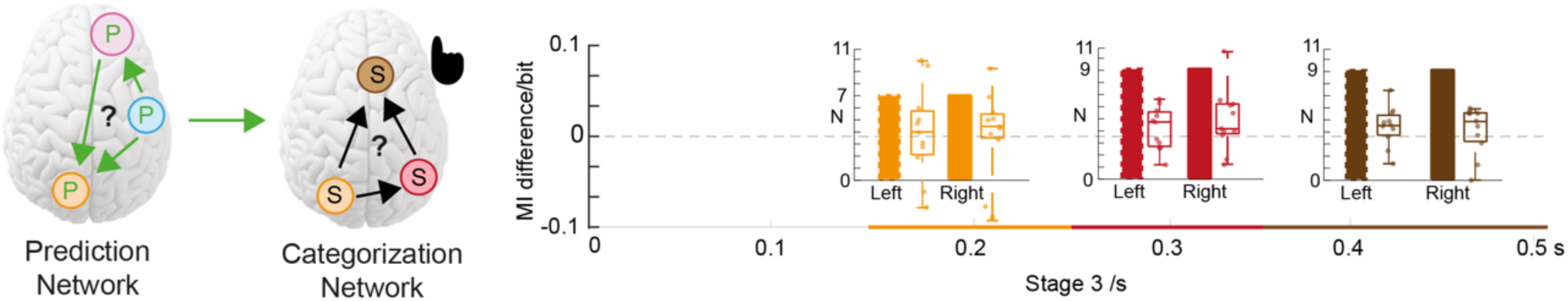
Prediction sharpens LSF vs. HSF Gabor discrimination across the categorization network. We compared representations of the same Gabor LSF vs. HSF discrimination–computed with MI(LSF vs. HSF Gabor; Stage 3 MEGt)–in predicted vs. non-predicted trials, separately for left- and right-located. On each occipital-ventral, parietal and pre-motor cortex source, valid predictions lead to significantly stronger representations of the LSF vs. HSF discrimination (i.e. peak MI, FWER, *p*<0.05, two-tailed) in the critical time window of the sequence (i.e., contra-lateral occipital-ventral: 150-250ms; parietal: 250-250ms; pre-motor: >350ms). Boxplots show peak MI difference between valid prediction vs. without prediction against the null hypothesis of no difference. Scatters show individual results. We replicated sharpening for left-(left boxplot and prevalence bar) and right-located trials (in right boxplot and prevalence bar), in 7/11 participants in contra-lateral occipital cortex, Bayesian population prevalence = 0.64 [0.33 0.85] (MAP [95% HPDI]), in 9/11 participants in parietal lobe and premotor cortex, Bayesian population prevalence = 0.81 [0.53 0.96], MAP [95% HPDI]. Besides, 2/11, 1/11 and 1/11 participants suggest decreased SF representations with prediction (i.e., dampening effect) in contra-lateral occipital, parietal, and premotor cortex.

In sum, Stage 2 LSF vs. HSF predictions on contra-lateral occipital sources preview LSF vs. HSF representations in the Prediction Network, sharpening Stage 3 stimulus representations which in turn speed up RTs in the Categorization Network.

## Discussion

We studied the network mechanisms that dynamically predict visual contents and then how, in turn prediction changes stimulus representation to facilitate categorization behavior. Our three-stage experimental design used a spatial cue to predict the left vs. right visual field location of an upcoming Gabor stimulus at Stage 1, to study the effects of prediction on stimulus representation specifically in the occipital cortex contra-lateral to stimulus presentation. The Stage 1 spatial cue was followed at Stage 2 by an auditory cue that predicted the LSF vs. HSF contents of the upcoming Gabor stimulus that appeared on the screen at Stage 3. We reconstructed a Prediction Network that propagates the auditory predictive cue from temporal (90-120ms post Stage 2 cue) to occipital cortex (200-400ms), via frontal cortex (120-200ms), all pre-stimulus. We showed that frontal cortex (PFC, in the dIPFC region) mediates the communication of the predictive cue from temporal to the left vs. right occipital cortex, depending on the cued location at Stage 1. When the Gabor stimulus is finally shown at Stage 3, we reconstructed post-stimulus Categorization Network that propagates the LSF vs. HSF feature from occipital-ventral cortex (150-250ms post Stage 3 Gabor), parietal lobe (250-350ms post Stage 3 Gabor) and premotor cortex (>350ms post Stage 3 Gabor). We then show how the Prediction Network changes the Categorization network and found that predictions sharpen LSF vs. HSF representations of the shown stimulus, from occipital cortex to pre-motor cortex, leading to faster categorization behavior. Together, our results reveal network mechanisms that communicate top-down the prediction of a specific content to occipital cortex, which sharpens the bottom-up representation of these contents in the stimulus to speed up behavior.

### Functional networks predict and then represent stimulus contents

Methodologically, we reconstructed a functional network that dynamically propagates a specific auditory prediction of visual contents (LSF vs. HSF) from temporal to occipital cortex, with mediation of the PFC. That is, PFC is necessary to propagate the predictive cue. Such connectivity analyses involve individual MEG sources acting as sending and receiving network nodes. Importantly, DFI functional connectivity differs from other signal-to-signal connectivity analyses (such as Granger causality or transfer entropy) because DFI isolates what the communication is about (at Stage 2, the auditory prediction of LSF vs. HSF) as a percentage of the full signal-to-signal connectivity ^21^. A similar logic isolated the mediatory role of PFC. Thus, DFI addressed the first question schematized in Figure 1, of the functional network of regions that dynamically (and multi-modally) propagate a prediction of visual information to the PFC that translates a prediction from auditory cortex into a predictive signal in occipital cortex that subsequently influences the representation of stimulus contents, when shown.

As the design auditory and visual inputs, one could object that Stage 2 effects merely reflect the auditory signals (not the predicted visual contents) spreading across the network. Our demonstration that PFC mediates the propagation addresses this objection by showing the high-level modulation which is distinct from the dynamic representation of the tone itself (as tested with the localizer prior to the experiment, see Supplemental Figure S3). Also, we proved the visual specificity of the Stage 2 prediction with end point in occipital cortex contra-lateral to predicted location (cf. Figure 3) and the representational overlap with the actual visual feature (see Supplemental Figure 9). Thus, the propagation of the visual prediction at Stage 2 is distinct from that of the auditory input.

To address the question of how Stage 2 prediction influences Stage 3 processing of the stimulus, we compared Stage 3 stimulus representation with and without prediction. A key unresolved question about the role of predictions is whether they sharpen vs. dampen stimulus representation ^1^. Evidence for one or the other typically relies on enhanced vs. impaired decoding performance of the predicted stimulus in the regions of interest ^7,14,16,17^. Here, we showed that predictions enhance the representation of LSF vs. HSF stimulus contents, locating these enhancements in source space and time. Most participants (7/11) showed that prediction sharpen LSF vs. HSF discrimination in occipital cortex and (9/11) in parietal cortex and (9/11) in premotor cortex, the latter relating to faster behavioral categorization. Thus, our evidence supports the hypothesis that the Prediction Network sharpens stimulus representation in the Categorization Network.

### The modulatory role of frontal cortex

An interesting finding of our functional network is that frontal cortex modulates the temporal to occipital communication of the predictive cue. More precisely, we located the sources with highest representation of the predictive cue in the dorsolateral prefrontal cortex (dlPFC, ^24^), often related to working memory ^25–27^, selective attention ^28^ and task performance ^29^. Frontal cortex could orchestrate the information of the auditory cue (i.e. upcoming LSF vs. HSF) together with the memory of the upcoming stimulus location (i.e. left vs. right visual field) and selectively prepare the contra-lateral occipital sources to the upcoming contents. Our results are compatible with this hypothesis, because representation of the prediction on occipital sources at Stage 2 is indeed contralateral to the predicted visual field where the stimulus will appear at Stage 3—i.e. left occipital sources for a predicted right visual field stimulus and vice versa. Future work that fuses MEG and high-field fMRI will seek to resolve the specific cortical laminar layer that receives the prediction at Stage 2 (e.g. central laminar layer ^30^), and how this prediction then interacts with the cortical layer representation of the feedforward flow when the stimulus is shown at Stage 3 (e.g. in peripheral laminar layers ^30^).

### Predictions and representations of face, object, body and scene stimuli

We used DFI to reconstruct the dynamic Prediction and Categorization Networks. Generalizing from Gabor stimuli to more naturalistic face, object and scene categorization tasks will incur several challenges that we can frame in the general context of studying the visual features that categorizes faces, objects and scenes _31,32_. A key challenge is that the stimulus features participants use to predict and then categorize real-world faces, objects and scenes, can differ across behaviors and levels of expertise (e.g. categorizing the same picture as “city” vs. “New York”) ^33,34^. We therefore need to characterize these features per participant and categorization task to then study their predictions and representations for behavior in functional networks ^35–41^. A methodological challenge will be to study the compositionality of visual predictions along the visual hierarchy, as they decompose from their integrated representation high in the visual hierarchy (e.g., right fusiform gyrus), to their contra-lateral components for occipital cortex, down to their simplest Gabor representation in the lower hierarchical levels. This will require fusion of brain measures such as high-field fMRI that finely tap into the laminar layers of these regions ^42^, with measures such as E/MEG ^43^ to trace the dynamics of these representations across layers in the occipito-ventral-dorsal streams.

Thus, to understand dynamic face, object and scene predictions and representations in the brain, we must understand the categorization task (e.g. “city vs. New York”), the hierarchical composition of features that represent each category in the participant’s memory, trace their hierarchical predictions in the feedback flow ^5^ and their subsequent representation in the feedforward flow when the stimulus is shown. This raises the thorny question of the granularity of features that can be predicted from memory at different hierarchical levels (in the visual and other modalities ^44,45^). In principle, once the compositionality of representations is understood, we could study how sensory hierarchies decompose their predictions from to facilitate stimulus processing and behavior.

## Conclusions

We sought to understand the propagation of specific predictions in a Prediction Network propagate and then how these predictions change the Categorization Network that processes the predicted contents in the stimulus. We showed that the Prediction Network dynamically propagates predictions of visual contents from temporal to occipital regions, via the modulatory role of frontal regions. Then, we showed that predicted contents were more sharply represented when the stimulus is shown in the Categorization Network, from occipital-ventral to pre-motor cortex, via parietal cortex, leading to faster decision behavior. Our Prediction and Categorization Networks in principle generalize to other stimulus features and sensory modalities.

## Methods

### Participants

Eleven participants (18-35 years old, mean=26.8, SD=3.0) took part in the experiment and provided informed consent. All had normal or corrected-to-normal vision and reported no history of any psychological, psychiatric, or neurological condition that might affect visual or auditory perception. The University of Glasgow College of Science and Engineering Ethics Committee approved the experiment (Application Number: 300210118).

### Stimuli

Stage 1 of the experimental design (see Figure 2) used two location cues (one for left- and one for right-located trials). Stage 2 used 3 different sweeping sounds, serving as LSF, HSF and neutral auditory cues. Stage 3 used 2 locations × 2 spatial frequencies × 3 orientations Gabor patches as stimuli. We detail them below.

#### Stage 1 Location Cues

Participants sat at a 182 cm viewing distance from the screen. We presented a green dot of 1 deg of visual angle diameter for 100 ms to the left (vs. right) of a fixation cross (2 deg of visual angle eccentricity).

#### Stage 2 SF Cues

Three 250 ms sweeping sounds started with auditory frequency of 196Hz (cueing LSF), 2217Hz (cueing HSF) or 622Hz (no prediction), with a sweep rate of 0.5 rising octave/second.

#### Stage 3 Gabor Stimuli

Left (vs. right)-presented Gabor patches were presented (diameter, 7.5 visual degrees; left and right eccentricity, 12.5 visual degree), with LSF (vs. HSF) contents of 0.5 cycle/degree (vs. 1.2 cycle/degree) shown at one of three randomly chosen orientations (−15 deg, 0 deg, +15 deg). Prior to the task, we calibrated the LSF and HSF Gabor contrast independently for each participant, using an adaptive staircase procedure (target accuracy set at 90%). On each calibration trial, a left (vs. right) green dot presented for 500ms predicted the upcoming left vs. right location of the LSF or (HSF) Gabor patch, itself presented for 100ms. Participant responded “LSF” vs. “HSF” vs. “don’t know” without feedback. We adaptively adjusted LSF vs. HSF contrast as follows:

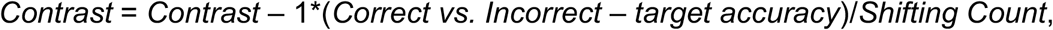

where *Shifting Count* counts the number of direction changes (i.e., increasing to decreasing, or decreasing to increasing). The adaptive staircase stopped when the adjustment step was < 0.01, setting each SF contrast for this participant’s Gabor in the actual experiment (Supplemental Table S4 provides each participant’s contrast values).

### Procedure

Each three-stage trial started with a central fixation cross presented for 500ms (Figure 1A accompanies the description below):

#### Stage 1

A green dot presented for 100ms appeared to the left or right of the central fixation cross, predicting the left vs. right location of the upcoming Gabor with a validity of 1. This was followed by a jittered blank screen [1000-1500ms].

#### Stage 2

Three sweeping sounds presented for 250ms predicted the Gabor stimulus presented at Stage 3. The 196Hz (vs. 2217Hz) sound predicted the upcoming LSF (vs. HSF) Gabor (both with .9 validity). The 622Hz sound was a neutral cue without predictive value. This neutral cue was followed by LSF vs. HSF Gabors with .5 probability, on 33% of the trials.

#### Stage 3

The LSF vs. HSF Gabor stimulus appeared at one of the three rotations on the left vs. right screen location for 100ms. The Gabor was either LSF or HSF, with one of three randomly chosen orientations, followed by a 750 to 1,250ms inter-trial interval (ITI) with jitter. We instructed participants to respond “LSF” vs. “HSF” vs. “Don’t know” as quickly and as accurately as they possibly could. They did not receive feedback.

The experiment comprised several blocks of 54 such trials (see Supplemental Table S5 for details). Participants performed 10-14 blocks in a single day, with short break between blocks. They completed the total of 38-45 blocks over 3-4 days. Participants completed at least 499 trials in each condition (of left vs. right presentation of LSF vs. HSF Gabors). Participants learned the correct relationships between the auditory cues and predicted SF within ∼2 blocks of trials, without explicit instructions. We therefore removed these first two blocks from all subsequent analyses.

#### Auditory localizer

Prior to the experiment, we ran an MEG localizer to model the bottom-up processing of each one of 3 auditory cues; For each cue, each localizer trial started with a blank screen for 500ms, followed by the auditory tone for 250ms, then a blank screen for 1250ms ITI. In a block of 12 trials, 10 of the trials presented the same tone and the two other tones were catch tones. Participants had to press a key whenever the tone was a catch tone. Each participant completed 36 such blocks (i.e., 12 blocks per type of tones), with block order of “low frequency”, “middle frequency”, “high frequency”, repeated 12 times.

### MEG Data Acquisition and Pre-processing

We measured participants’ MEG activity with a 248-magnetometer whole-head system (MAGNES 3600WH, 4-D Neuroimaging) at a 508Hz sampling rate. We performed the analysis according to recommended guidelines using the FieldTrip toolbox ^46^ and in-house MATLAB code.

For each participant, we epoched the raw data into trial windows, separately for each stage: Stage 1, -200ms pre-dot onset to 1,000ms post-dot onset (henceforth [-200ms 1,000ms] around onset); Stage 2: [-200ms 1,000ms] around sweeping sound onset; Stage 3: [-200ms 600ms] around Gabor patch onset. Then, we applied 1Hz high-pass filter (5^th^ order two-pass Butterworth IIR filter) to the epoched data, removed the line noise using discrete Fourier transform and de-noised via a PCA projection of the reference channels. We rejected noisy channels with a visual selection and rejected jump and muscle artifacts with automatic detection. We decomposed the output dataset with ICA, identified and removed the independent components corresponding to artifacts (eye movements, heartbeat—i.e. 2-4 components per participant).

We then resampled the output data at 512 Hz, low-pass filtered the data at 25Hz (5^th^ order Butterworth IIR filter), specified the time of interest between 0-500ms (post cue at Stage 2; Gabor stimulus at Stage 3), and performed the Linearly Constrained Minimum Variance Beamforming (LCMV) analysis to reconstruct the time series of sources on a 6mm uniform grid warped to standardized MNI coordinate space.

Following the above steps, for each participant we obtained single-trial time series of 12,773 MEG sources at a 512Hz sampling rate between 0 and 500ms that we used to analyze the dynamic information processing in the Prediction and Categorization Networks—i.e. at Stages 2 and 3, see Figure 1. Supplemental Table S6 reports the numbers of participant trials that remained after pre-processing and LCMV analysis.

#### Auditory Localizer

We applied the same pre-processing pipeline to the MEG localizer, using the epoched data [-200ms 500ms] around tone onset. We applied the LCMV analysis 0-500ms post tone, to reconstruct the source representation of the MEG localizer data. Supplemental Table S6 summarizes the numbers of trials that remained after pre-processing and LCMV analysis.

## Analyses

### Cueing improves behavior

At a group-level, we discarded invalid trials and applied a 2 (left vs. right location cues) × 2 (valid informative vs. neutral SF cues) × 2 (LSF vs. HSF Gabor patches) ANOVA on the median RTs (excluding incorrect response and outliers) and on the accuracy of valid trials of all participants. We found a significant main effect of valid informative vs. neutral SF cueing on RTs, showing that predicted trials are significantly faster than non-predicted trials (*F(1,86)*=20.8, *p*=0.001); and a significant interaction effect between location cue and Gabor SF (*F(1,86)*=17.4, *p*=0.002). Further analysis showed that this informative vs. neutral cueing effect is significant (*p*<0.05, after Bonferroni correction) for each of the 4 experimental conditions (left vs. right locations × low vs. high SFs), quantified by a paired-sample t-test, independently for each condition. For categorization accuracy (ACC), the ANOVA was significant only for valid informative vs. neutral cueing, showing that ACC is significantly higher in in predicted than non-predicted trials (*F(1,86)*=22.5, *p*=0.0008); and a significant interaction between location cue and Gabor SF (*F(1,86)*=13.8, *p*=0.004). Further analysis showed that this effect of SF cue is significant (*p*<0.05, Bonferroni correction) for all but the left-LSF experimental conditions (paired-sample t-test independent for each condition).

We analyzed the RTs (excluding trials with incorrect responses and outlier trials) and ACC of each individual participant, replication in each the significant main effect of SF cue on RTs (tested with the 2 × 2 × 2 ANOVA) and also examined the effect of SF cue for experimental conditions (independent-sample *t*-tests). Supplemental Table S2 summarizes these ANOVA and independent-sample *t*-tests on RT statistics. Supplemental Table S3 summarizes the statistics of 2 × 2 × 2 ANOVA and chi-squared test for the accuracy data.

### Stage 2: Prediction Network

#### Prediction representations

To understand the Stage 2 network of regions that propagates the LSF vs. HSF auditory prediction prior to stimulus onset, we computed the representation of the cue across the whole brain, separately for left- and right-located trials.

For each participant, we computed the single-trial MI(<LSF vs. HSF auditory cue; Stage 2 MEG_t_>), at each time point from 0 to 400ms following Stage 2 auditory cue onset, on each occipital source (lingual gyrus, cuneus, inferior occipital gyrus), temporal (fusiform gyrus, inferior temporal gyrus, middle temporal gyrus, superior temporal gyrus), parietal (superior parietal lobe, inferior parietal lobe, angular gyrus, supramarginal gyrus), premotor (precentral gyrus, postcentral gyrus), and frontal (orbitofrontal gyrus, inferior frontal gyrus, middle frontal gyrus, medial frontal gyrus, superior frontal gyrus).

#### Prediction periods clustering

To compute the number of space x time periods of prediction representations, we applied k-means clustering analysis on all 4413 × 204 (source x time points) dimensional trials as follows:

*Step 1: Peak time extraction*. First, for each participant, and independently for left- and right-located trials and source, we extracted the peak time MI(<LSF vs. HSF auditory cue; Stage 2 MEG_t_>), 0 and 400ms post auditory cue onset.

*Step 2: Matrix computation*. Across participants and condition, we counted the numbers of sources per ROI (occipital, temporal, parietal, pre-motor and frontal) that peak during each 10ms-step time window between 0 and 400ms post auditory cue onset (i.e. 39 time windows), producing a ROI x time matrix of MI peaks.

*Step 3: Clustering*. We k-means clustered (k = 1 to 30, repeating 1,000 times) the matrix from Step 2, using the 39 time windows as samples and selected k as the elbow of the within-cluster sums of point-to-centroid distances metric.

Supplemental Figure S1 shows Stage 2, with k = 4 as a good solution, starting with a period 0, before any prediction representation, and then 3 distinct timed periods with temporal, frontal, occipital of peak representations of the prediction.

#### Prediction network nodes (supports Figure 3A)

To reveal the dynamics of MI(<LSF vs. HSF auditory cue; Stage 2 MEG>) representation of the prediciton, we localized the source peaking around the first peak in the 90-120ms time window (start), the last peak in 120-200ms (midway) and >200ms (end). We computed the group mean of these 3 source-localized peaks across participants (see Figure 3A for group mean and Supplemental Figure S2 for each individual result).

#### Prediction network reconstruction (supports Figure 3B)

To reconstruct the Stage 2 Prediction Network, we computed Directed Feature Information (DFI, where F is the auditory cue predicting the upcoming LSF vs. HSF Gabor) in each participant, for each pair of identified network nodes (i.e., sender: temporal, receiver: frontal; sender: frontal, receiver: occipital) as follows:

*Step 1: Source selection*. We selected the highest MI source for the sending and receiving regions in the time window of interest (temporal: 90-120ms, frontal: 120-200ms, occipital: >200ms).

*Step 2: Directed Information (DI)*. DI (i.e. Transfer Entropy) quantifies all the information communicated from sending to receiving sources, removing information sent from the receiver itself. For the receiver at time *x*, with a communication delay *y* from the sender, DI is computed as the Conditional Mutual Information (CMI) between *RA*_*x*_ and *SA*_*x-y*_ conditioned on *RA*_*x-y*_:

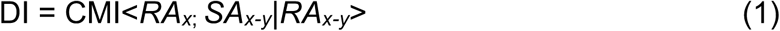

Thus, we computed DI between each sender and receiver source, for each receiver time point between 0 and 400ms post auditory cue onset, and for each communication delay between 0 and 300ms. This produced the receiver-time × transfer-delay DI matrix. Supplemental Figure S4A shows the DI computation.

*Step 3: DI conditioned on Feature (DI*|*F)*. DI|F removes from DI the information communicated about the predictive LSF vs. HSF feature itself. We computed DI|F for each receiving-time × communication-delay and show it in the Supplemental Figure S4B.

*Step 4: DFI*. The difference between DI and DI|F isolates the information communicated about the predictive cue. We computed DFI as:

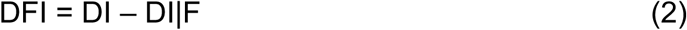

for each receiving-time × communication-delay cell of the matrix and show it in the Supplemental Figure S4C.

*Step 5: Statistical significance*. We repeated 200 times DFI computations with shuffled feature labels (i.e., LSF vs. HSF), using as statistical threshold the 95^th^ percentile of the distribution of 200 maxima (each taken across the DFI matrix of each shuffled repetition, FWER, *p*<0.05, one-tailed).

We applied Step 1-5 to reconstruct the Stage 2 Prediction Network of each individual participant. Supplemental Figure S5 shows the individual participants DFI networks; Figure 3B shows the group average network.

#### Prediction network modulation (supports Figure 4)

We then tested whether frontal cortex is a necessary mediator of Stage 2 prediction communications between temporal and occipital cortex, by isolating the role of the frontal region in these communications. We then compared network communications with and without frontal mediation. The steps below detail how we computed frontal mediation in the Prediction Network of each participant.

*Step 1: Frontal Mediation, DFI*. On the selected temporal and occipital sources, for receiving time points between 0 and 400ms post auditory cue onset and for each delay between 0 and 300ms, we computed the receiving-time × communication-delay of temporal-to-frontal DFI and then frontal-to-occipital DFI (each computed as in <u>*Prediction network reconstruction*</u>). This quantifies the mediating role of the frontal region in the communication of the predictive cue (cf. Figure 4B).

*Step 2: Direct Communication, DFI*|*Frontal*. To isolate the role of frontal mediation, we also computed temporal-to-occipital DFI conditioned on the frontal activity. Specifically, for each time point in the combination of (1) receiving time *x* between 0 and 400ms post auditory cue onset (2) communication delay *y* between 0 and 300ms (3) and mediation time *z* between receiving time and sending time (i.e., *x* and *x-y*), we computed DFI received by occipital at time *x*, sent by temporal at *x-y*, conditioned on frontal activity at time *z*.

This produced the 3D DFI receiving-time × communication-delay × mediation-time conditioned DFI matrix. We took the minimum conditioned DFI across the mediation time as the directed communication (i.e., without frontal mediation, Figure 4A).

*Step 3: Statistical significance*. We recomputed Steps 1 and 2 and their difference, shuffling the LSF vs. HSF labels—i.e. 200 repetitions, using the 95^th^ percentile of 200 maxima as statistical threshold, each maximum taken across the DFI minus DFI|F matrix of each shuffled repetition, FWER, *p*<0.05, one-tailed. This isolated the receiving-time × communication-delays showing significant enhancement with vs. without frontal mediation.

We applied Steps 1-3 to each participant. Figure 4A and B show the results of a typical participant. Supplemental Figure S6 shows all individual results. Figure 4C shows the group mean difference and its Bayesian prevalence.

### Stage 3: Categorization Network

#### Stimulus representations

To reconstruct the Stage 3 Categorization Network, we computed for each participant the dynamics of LSF vs. HSF Gabor stimulus representation across the whole brain, separately for left- and right-located trials—i.e. MI(LSF vs. HSF Gabor; Stage 3 MEG_t_), on each source in occipital, temporal, parietal, premotor and frontal regions, at each time point from 0 to 500ms following Gabor onset.

#### Categorization periods clustering

To compute the number of space x time stimulus representations period, we applied again k-means cross-trials clustering analysis on all 4,413 sources x 256 time points as follows:

*Step 1: Peak time extraction*. First, for each participant, and independently for left- and right-located trials and source, we extracted the peak LSF vs. HSF representation MI in 50 10-ms time windows spanning 0-500ms post Gabor.

*Step 2: Matrix computation*. Across participants and conditions, we counted the numbers of sources per ROI (occipital, temporal, parietal, pre-motor and frontal) and time window the number of sources that peak, producing a ROI x time matrix of MI peaks.

*Step 3: Clustering*. We k-means clustered (k = 1 to 30, repeating 1,000 times) the matrix from Step 2, using the 50 time windows as samples and selected k as the elbow of the within-cluster sums of point-to-centroid distances metric.

Stage 3 comprised k = 4 clusters (see Supplemental Figure S1). A first period with no LSF vs. HSF stimulus representation, followed an occipital-ventral (150-250ms, start), parietal (250-350ms), and premotor-frontal (>350 ms) periods of stimulus representation.

#### *Categorization network nodes* (supports Figure 5A)

To reveal the dynamics of MI(LSF vs. HSF Gabor; Stage 3 MEG), in each participant, we localized the source peaking around in each one of the three representational periods. We then computed the group mean of these 3 sources across participants (Supplemental Figure S7 for each individual result). Figure 5A presents the group mean.

#### *Categorization network reconstruction* (supports Figure 5B)

To reconstruct the Stage 3 Categorization Network that communicates the Gabor SF across occipital, parietal, premotor regions identified earlier, we computed DFI communications of the LSF vs. HSF stimulus information. That is, in each participant, for each pair of regions (i.e., sender: occipital, receiver: parietal; sender: parietal, receiver: premotor), we performed the following three steps.

*Step 1: Source selection*. We selected one sender and one receive source with highest Stage MI representation of Gabor LSF vs. HSF in the time window of interest (occipital: 150-250ms, parietal: 250-350ms, premotor: >350ms).

*Step 2: DFI*. For each receiver time points between 0 and 500ms post Gabor stimulus onset, and for each sender delays between 0 and 300ms, we computed the receiver-time × communication-delay of LSF vs. HSF stimulus representation with DFI (see specific computations in <u>*Prediction network reconstruction*</u>).

*Step 3: Statistical significance* was established recomputing DFI with shuffled LSF vs. HSF labels—i.e. 200 repetitions, using as statistical threshold the 95^th^ percentile of 200 maxima, each taken across the DFI matrix of each shuffled repetition, FWER, *p*<0.05, one-tailed.

We applied Steps 1-3 in each participant, reconstructing the occipital-to-parietal and parietal-to-premotor network that communicates the LSF vs. HSF Gabor contents (Supplemental Figure S8 shows all individual results; Figure 5B shows the group average).

### Stage 2 to Stage 3: Interactions of Prediction Network on Categorization Network

#### Stage occipital prediction overlaps with Stage 3 LSF vs. HSF stimulus representation

To determine whether per-trial Gabor predictions at Stage 2 comprise the same SF contents as when Gabor stimuli are actually processed at Stage 3, we quantified the overlap of Gabor LSF vs. HSF representations between the two stages. We used information theoretic redundancy, quantified with co-information. Co-I quantifies the information about LSF vs HSF that is shared between or common to the Stage 2 and Stage 3 occipital MEG responses.

Step 1: *Source selection*. We selected one Stage 2 source (Period 3, >200ms) and one Stage 3 source (Period 4, 150-250ms) with highest MI in the occipital cortex contra-lateral to predicted Gabor presentation.

Step 2: *Co-Information*. We quantified the occipital representational overlap between Stage 2 Gabor LSF vs. HSF predictions and actual Stage 3 LSF vs. HSF Gabor representations as Co-I(<LSF vs. HSF, Stage 2 MEG_t_; Stage 3 MEG_t_>) on trials with valid predictions, separately for left- and right-localized Gabor stimuli.

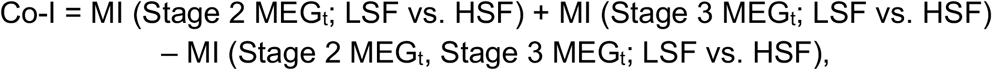

We computed Co-I every 2ms between 0 and 400ms Stage 2 post auditory cue onset and every 2ms between Stage 3 0 and 500ms post Gabor onset, producing a 2D Co-I matrix per participant (Stage 2 time × Stage 3 time).

Step 3: *Statistical significance* was established by recomputing Co-I with shuffled LSF vs. HSF labels, 200 repetitions, using as statistical threshold the 95^th^ percentile of 200 maxima (each taken across the 2D Co-I matrix of each shuffled repetition, FWER, *p*<0.05, one-tailed). To reflect when and how strongly Stage 2 stimulus prediction overlaps with its Stage 3 representation, for Stage 3 time point, we selected the maximum Stage 2 Co-I value.

We repeated the above Steps 1-3 for each participant.Supplemental Figure S9 shows the individual and group results.

#### Prediction modulates Categorization Network source activity and RT

To demonstrate where and when valid predictions modulate premotor MEG activity to facilitate behavior, we compared the effect of valid prediction vs. no prediction at Stage 2 on Stage 3 premotor brain activity and behavioral RT.

Step 1: *Source selection*. We selected one premotor cortex source at Stage 3, Period 6 with highest MI(<LSF vs. HSF Gabor; Stage 3 MEG_t_ >) >350ms post Gabor onset.

Step 2: *Co-Information*. We computed positive Co-I(<predicted vs. non-predicted; Stage 3 MEG_t_; RT>), information theoretic redundancy, as follows:

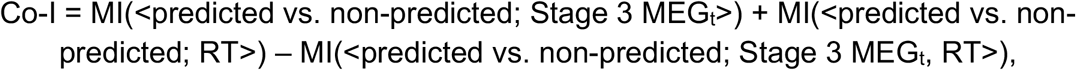

every 2ms between 0 and 500ms post Gabor onset, producing a vector in Stage 3 time.

Step 3: *Statistical significance* was established by recomputing the Co-I with shuffled LSF vs. HSF labels, 200 repetitions, using as statistical threshold the 95^th^ percentile of 200 maxima, each taken across the vector of each shuffled repetition, FWER, *p*<0.05, one-tailed.

We applied Step 1-3 to each participant.Supplemental Figure S9 shows the individual and group results.

#### Prediction sharpens stimulus representation (supports Figure6)

To understand how Stage 2 predictions of LSF vs. HSF facilitate Stage 3 their Stage 3 categorization when the stimulus is shown, we compared LSF vs. HSF Gabor representations between Stage 3 predicted and non-predicted trials, in each participant and Categorization Network region (i.e., contra-lateral occipital-ventral, parietal, premotor). Specifically, we computed MI as follows:

Step 1: *Source selection*. We selected one Stage 3 source per region with highest MI(LSF vs. HSF Gabor; Stage 3 MEG_t_) during the time window of interest (occipital-ventral: 150-250ms; parietal: 250-350ms; premotor: >350ms).

Step 2: *MI computation*. For each selected source, we computed source-by-time MI(LSF vs. HSF Gabor; Stage 3 MEG), every 2ms between 0 and 500ms post Gabor onset, separately for predicted and non-predicted trials. For this computation, we matched number of predicted trials with non-predicted trials (random selection). We averaged the MI matrices for predicted trials from 5 such random trial selections.

Step 3: *Statistical significance of difference* was established by recomputing the source-by-time MI with shuffled predicted and unpredicted trials (repeated 200 times), calculating the difference of peak between recomputed predicted and non-predicted MI in the time window of interest, using as statistical threshold the 95^th^ percentile of 200 maxima, each taken across the source-by-time difference of each shuffled repetition (FWER, *p*<0.05, two-tailed).

We repeated above Step 1-3 for each participant. Figure 6 shows the group-level results.

### Auditory localizer representation

To compare the spatio-temporal pattern that represents the predictive cue to that which merely represent the bottom-up auditory input without predicted contents, we computed in each participant the dynamics bottom-up representation of the different tones using localizer data—i.e as MI(low frequency vs. high frequency tone; localiser MEG_t_), between 0 to 400ms following tone onset, on each occipital, temporal, parietal, premotor, and frontal source.

## Supplemental Materials

**Supplemental Table S1.** Group-level effect of cueing on mean LSF vs. HSF, left- and right-located Gabor categorization RTs (paired samples t-tests).

| Gabor type | RT (predicted, ms) | RT (non-predicted, ms) | Improvement on RT (ms) | *t* value | *p* value |
| --- | --- | --- | --- | --- | --- |
| Left LSF | 530.9 | 456.8 | 74.1 | $t_{(10)} = 3.60$ | $p = 0.005$ |
| Left HSF | 555.4 | 447.7 | 107.7 | $t_{(10)} = 5.87$ | $p = 0.0002$ |
| Right LSF | 556.3 | 483.6 | 72.6 | $t_{(10)} = 3.37$ | $p = 0.007$ |
| Right HSF | 525.9 | 429.5 | 96.4 | $t_{(10)} = 4.82$ | $p = 0.0007$ |

**Supplemental Table S2.**
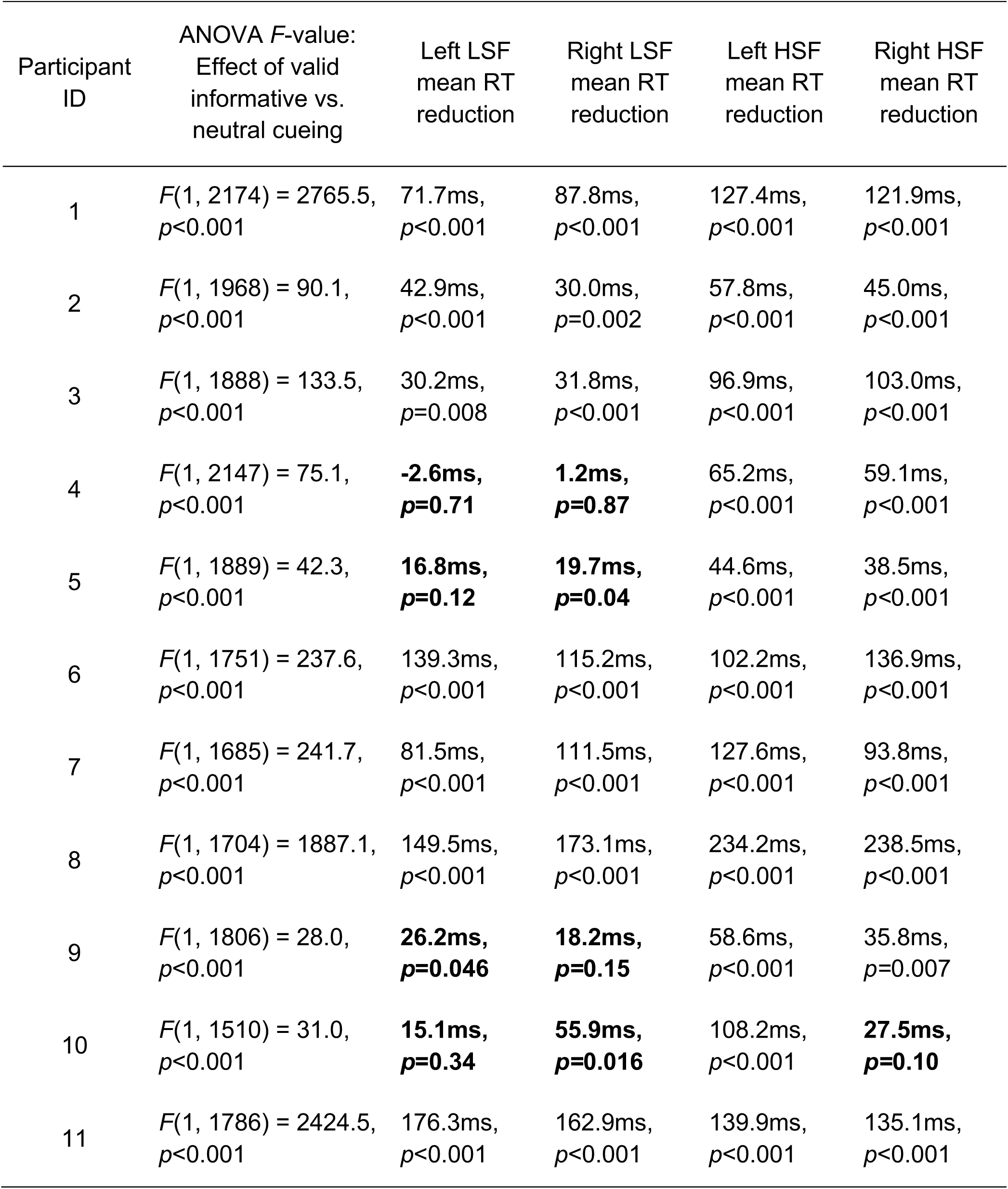
Individual-level effect of predicted vs. non-predicted trials on categorization RTs of Gabor patches—i.e. 2 (left vs. right-located) x 2 (LSF vs. HSF) ANOVA and independent samples *t*-test, before Bonferroni correction. **Bold** indicates **non-significant** results (*p*>0.05, Bonferroni-corrected).

**Supplemental Table S3.**
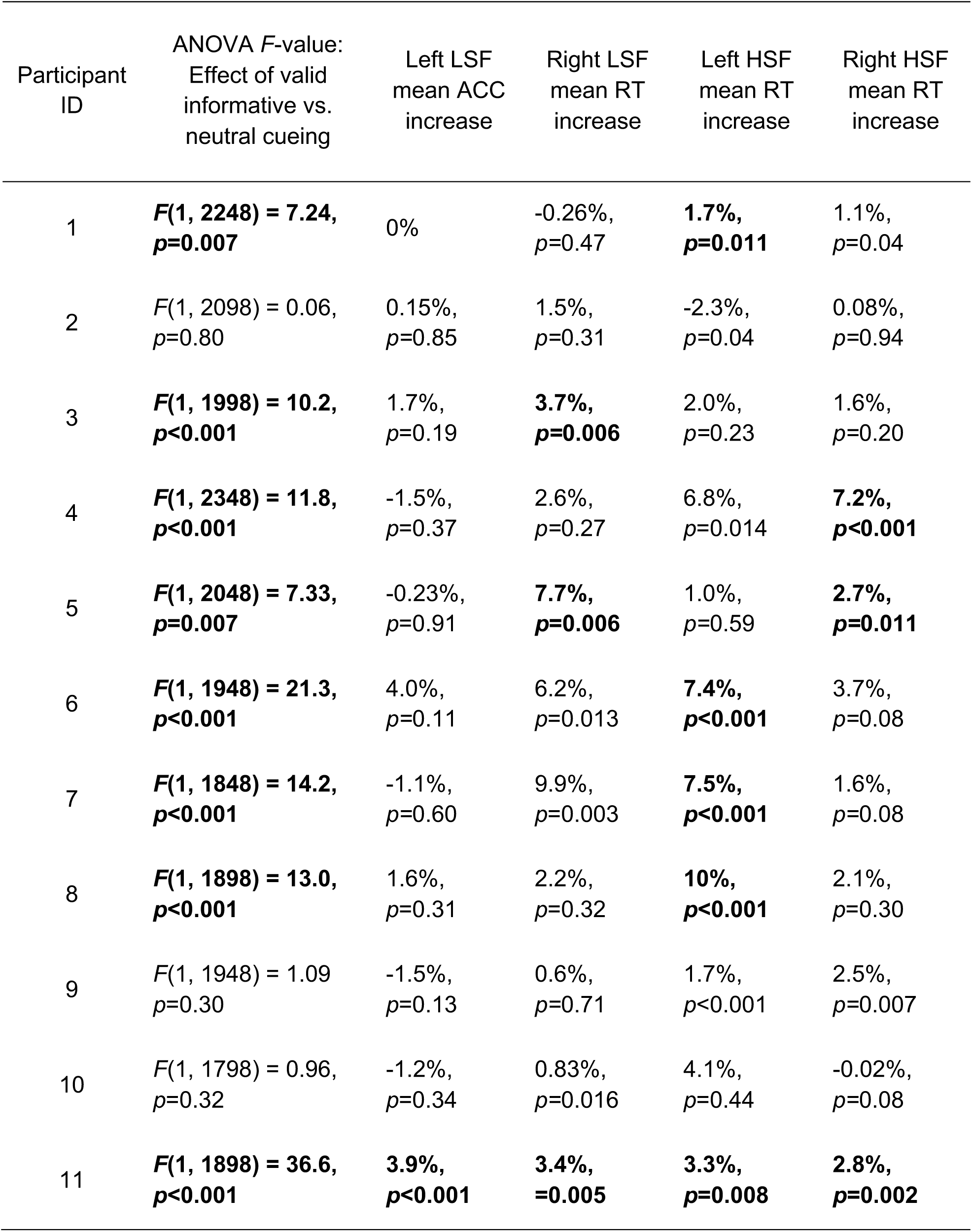
Individual-level effect of predicted vs. non-predicted on categorization accuracy of Gabor patches—i.e. 2 (left vs. right-located) x 2 (LSF vs. HSF) conditions, ANOVA and chi-squared tests, before Bonferroni correction. **Bold** indicates **significant** results (*p*<0.05, Bonferroni-corrected).

**Supplemental Table S4.**
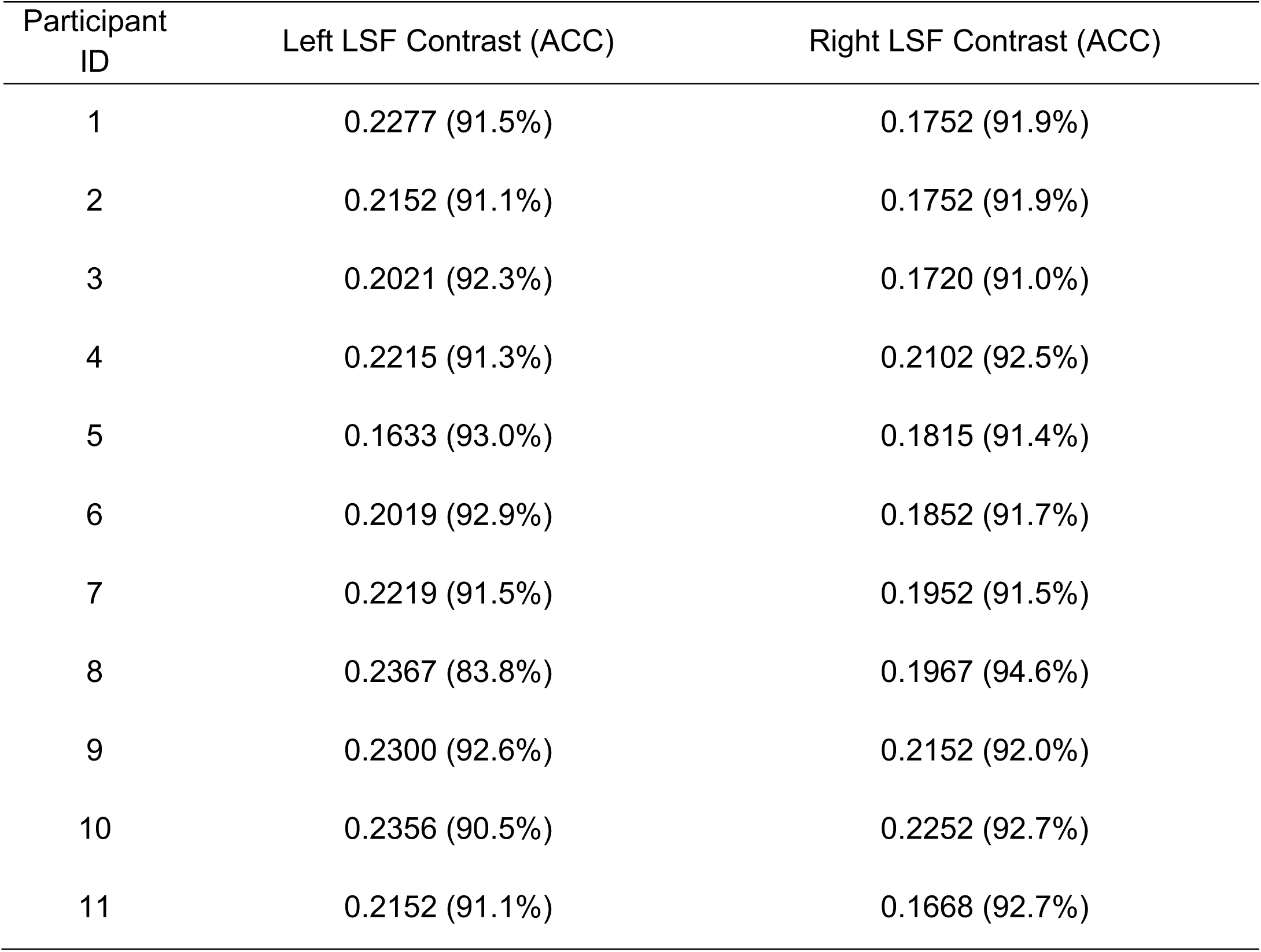
Individual-level contrast and accuracy of LSF vs. HSF Gabor patches in contrast threshold testing.

| Participant ID | Left LSF Contrast (ACC) | Right LSF Contrast (ACC) |
| --- | --- | --- |
| 1 | 0.2277 (91.5%) | 0.1752 (91.9%) |
| 2 | 0.2152 (91.1%) | 0.1752 (91.9%) |
| 3 | 0.2021 (92.3%) | 0.1720 (91.0%) |
| 4 | 0.2215 (91.3%) | 0.2102 (92.5%) |
| 5 | 0.1633 (93.0%) | 0.1815 (91.4%) |
| 6 | 0.2019 (92.9%) | 0.1852 (91.7%) |
| 7 | 0.2219 (91.5%) | 0.1952 (91.5%) |
| 8 | 0.2367 (83.8%) | 0.1967 (94.6%) |
| 9 | 0.2300 (92.6%) | 0.2152 (92.0%) |
| 10 | 0.2356 (90.5%) | 0.2252 (92.7%) |
| 11 | 0.2152 (91.1%) | 0.1668 (92.7%) |

**Supplemental Table S5.**
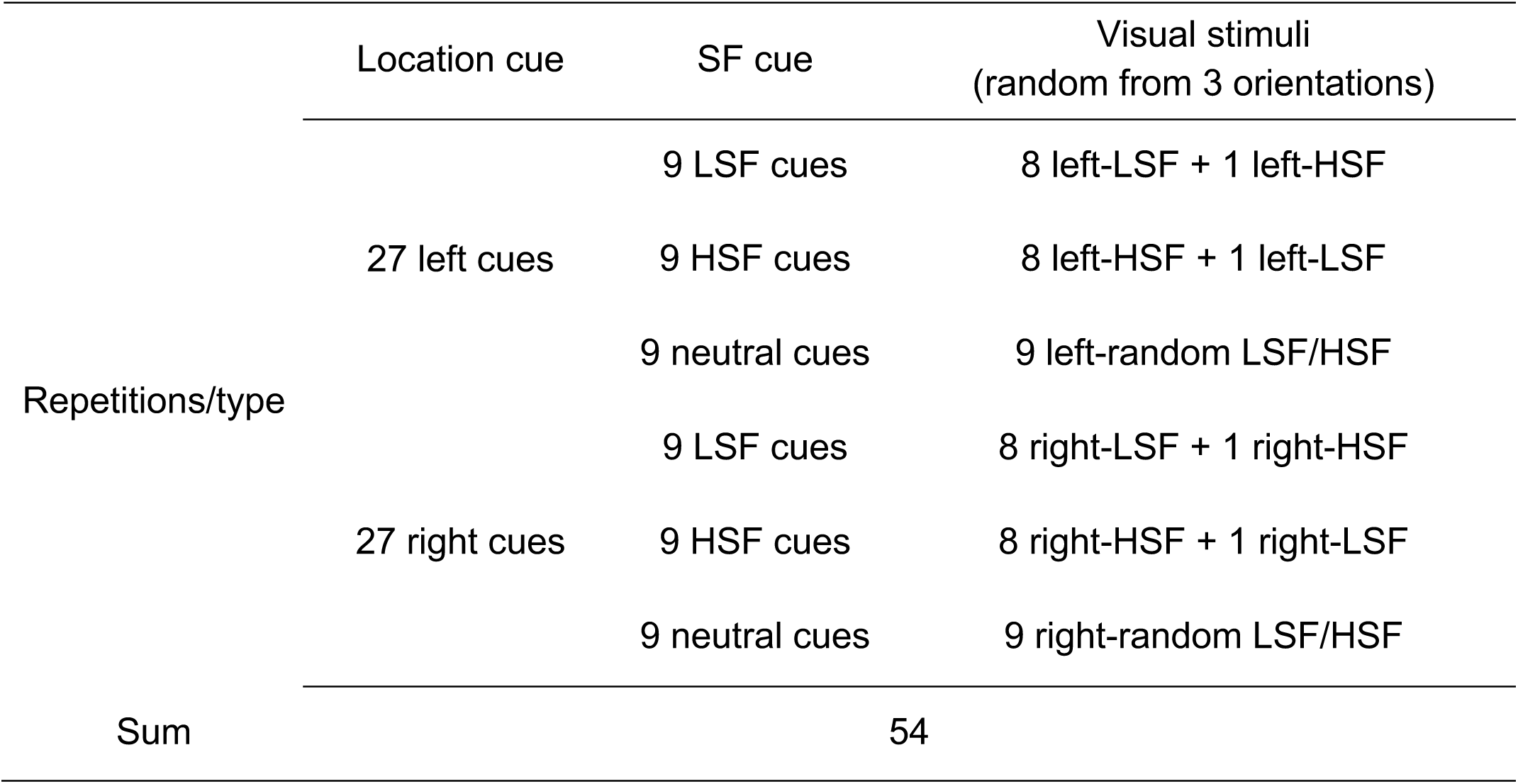
Stimulus repetition in one cueing-categorization block

**Supplemental Table S6.** Trials remaining following pre-processing and LCMV analysis

| Stage | Mean trial number/participant | Range |
| --- | --- | --- |
| Stage 1 | 1609 | 1492-1759 |
| Stage 2 | 1604 | 1505-1751 |
| Stage 3 | 1620 | 1507-1743 |

**Supplemental Figure S1.**
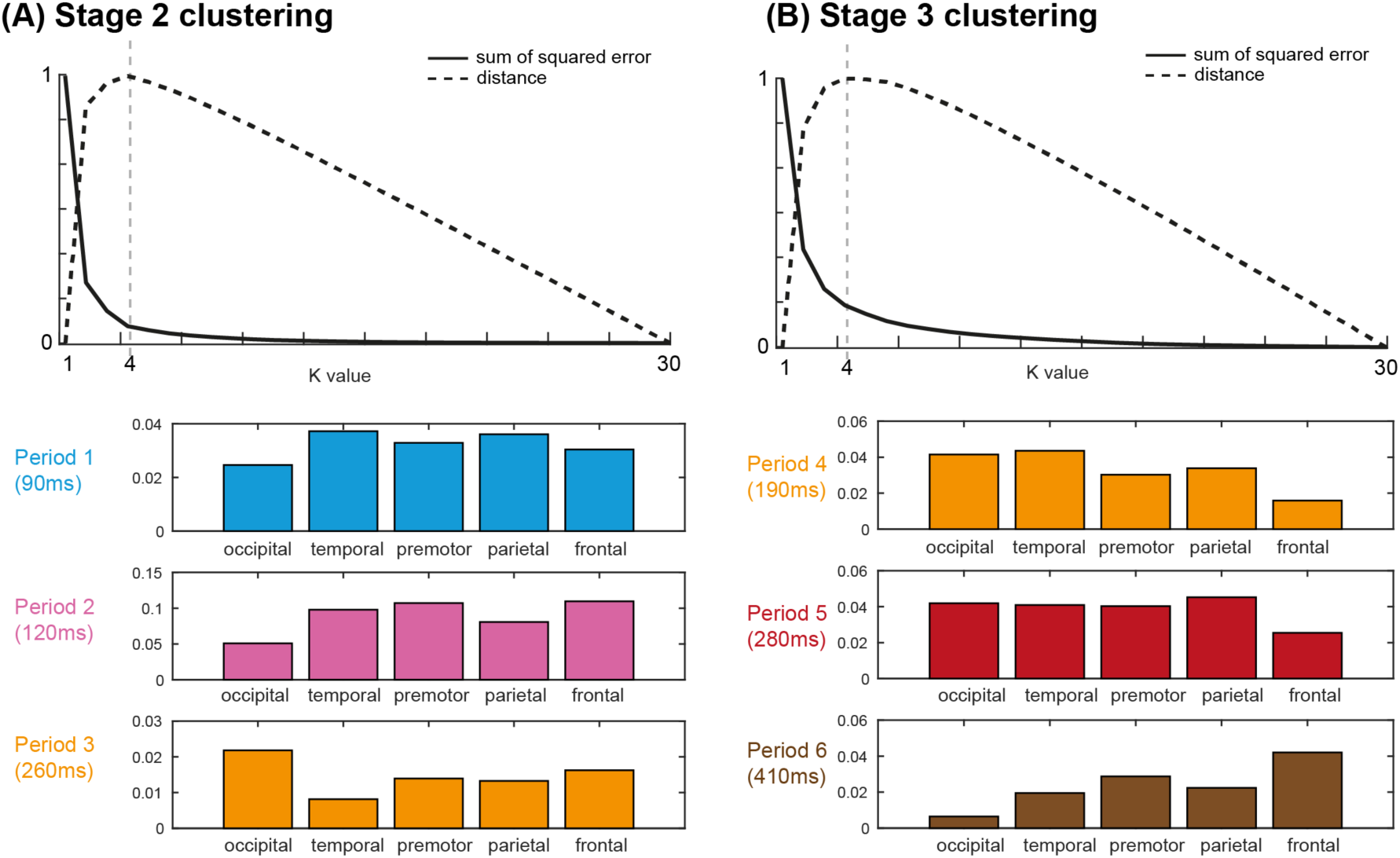
Clustering of representational time windows at (A) Stage2 (prediction representation) and (B) Stage 3 (stimulus representation). k-means analysis, with k = 1..30, with sum of squared error of each k (plain curve) and distance of the plain curve from straightline (dashed curve). We selected k with furthest distance (elbow method). Following a baseline period (Period 0), the bar plot shows the percentage of sources peaking at the center time of each period, at occipital, temporal, premotor, parietal, frontal respectively. Stage 2 comprises 3 periods of predictive cue representations, dominated by temporal, frontal and occipital. Stage 3 contains 3 periods of stimulus representations, dominated by occipital-temporal, parietal and premotor-frontal.

**Supplemental Figure S2.**
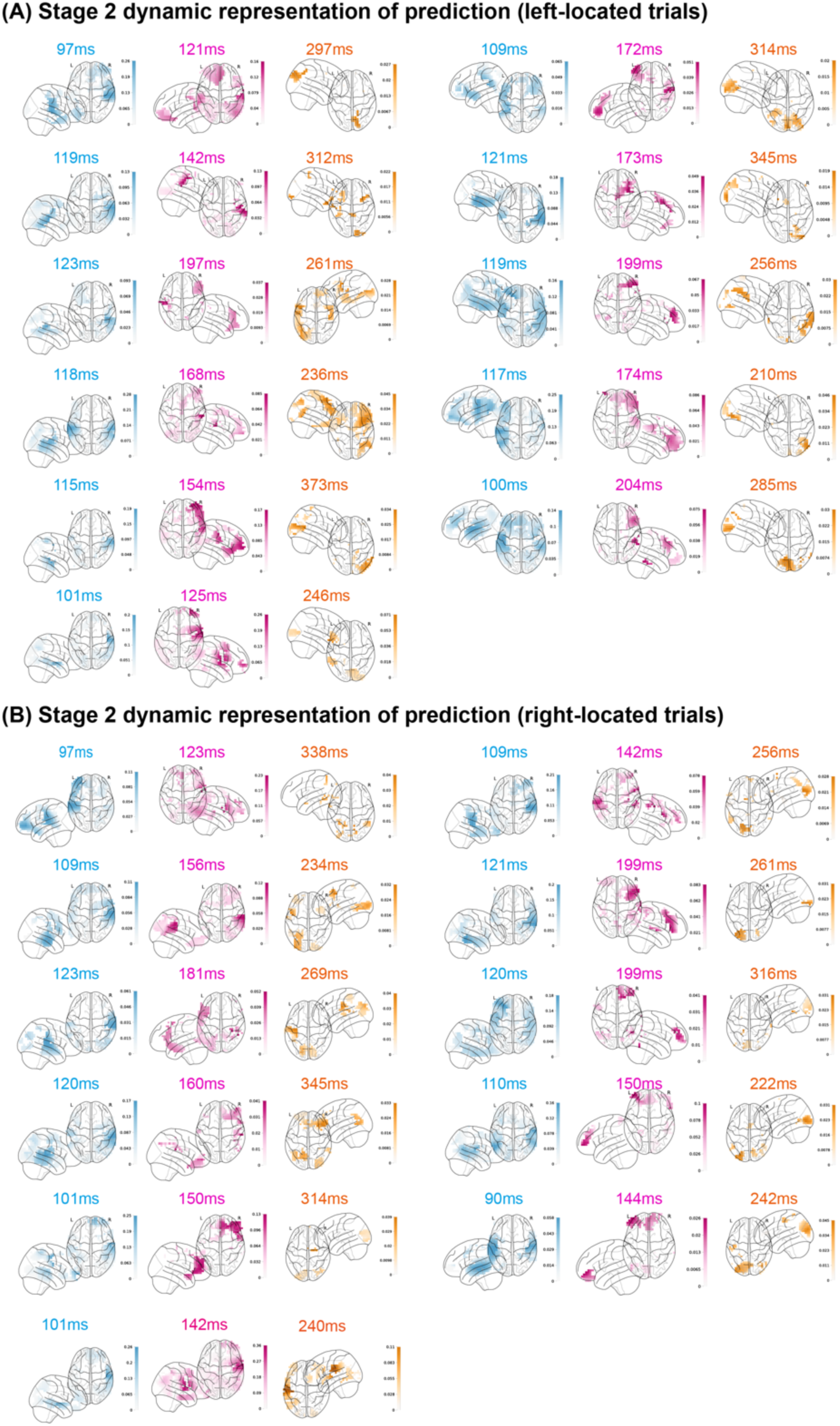
Stage 2: Dynamic representation of LSF vs. HSF prediction in individual participants. Separately for **(A) left-located trials** and **(B) right-located trials**, we computed the single-trial MI(LSF vs. HSF auditory cue; Stage 2 MEG_t_), at each time point from 0 to 400ms following auditory cue onset, on each of 4413 sources (see *Methods, Prediction representations*). We then extracted the peak MI time for each source. To reveal the representational dynamics, we localized the source with first MI peak in the 90-120ms time window (start, in blue), the last peak in the 120-200ms time window (midway, in pink) and peak in >200ms time window (end, in orange). The brain plots localize the sources with peaks in each time window.

**Supplemental Figure S3.**
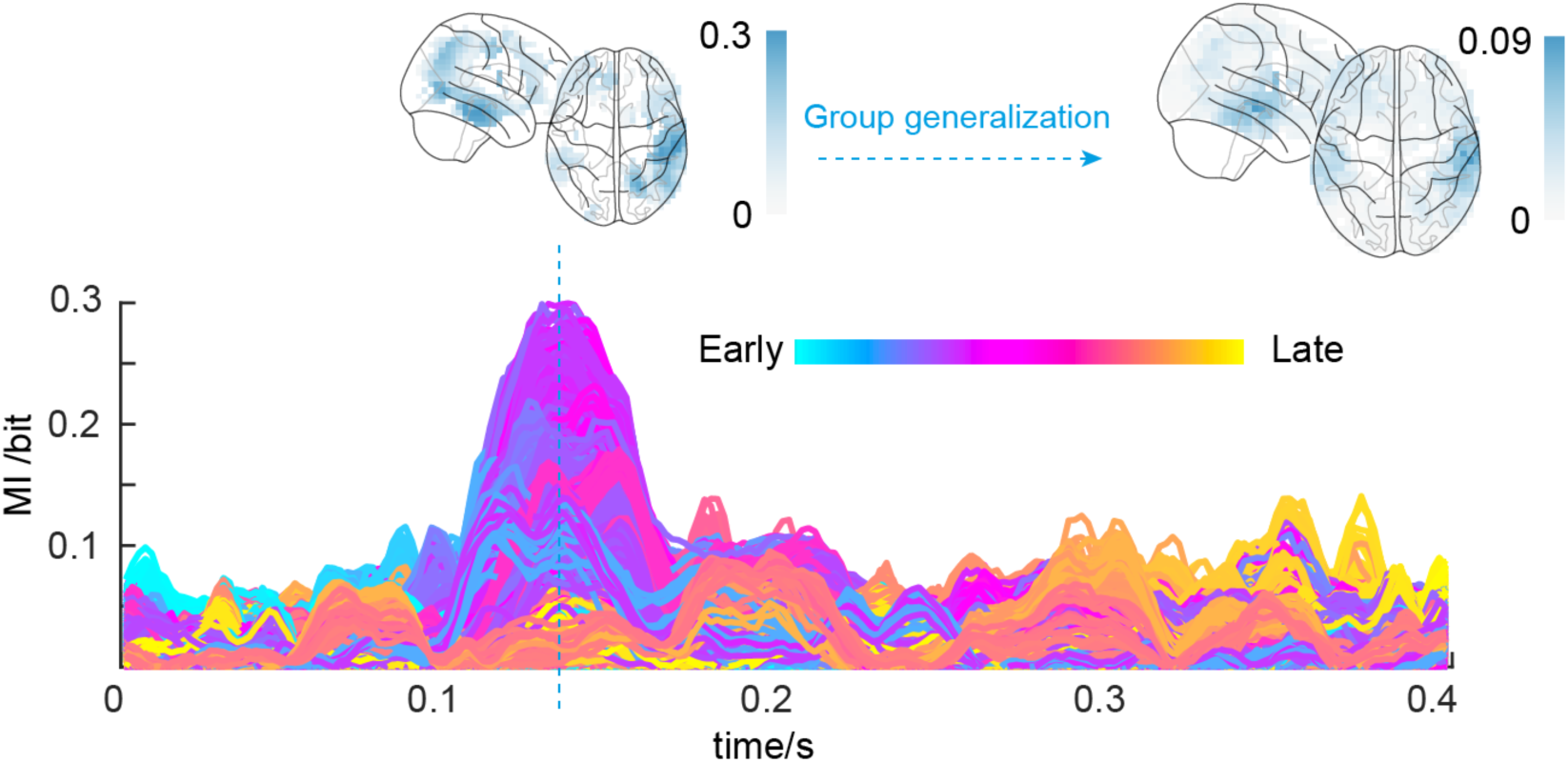
Tone representation in auditory localizer (run before the cueing experiment). For a typical participant, we computed the representation of the predictive tone (as MI(LSF vs. HSF cue; MEG_t_), Y-axis), between 0 and 0.4s post tone (X-axis), on each source in temporal (fusiform gyrus, inferior temporal gyrus, middle temporal gyrus, superior temporal gyrus), frontal (orbitofrontal gyrus, inferior frontal gyrus, middle frontal gyrus, medial frontal gyrus, superior frontal gyrus) and occipital regions (lingual gyrus, cuneus, inferior occipital gyrus). The curves show these per-source time courses of tone representation, color-coded by their (ranked) peak MI time. Glass brains below reveal that the localized sources peak in the only period—i.e. [122-142ms], showing that representation of the tone in the localizer (i.e. in the absence of a prediction) only activates the temporal lobe (cyan). The right glass brains show the average of this analysis across participants, revealing the same consistent result at group-level.

**Supplemental Figure S4.**
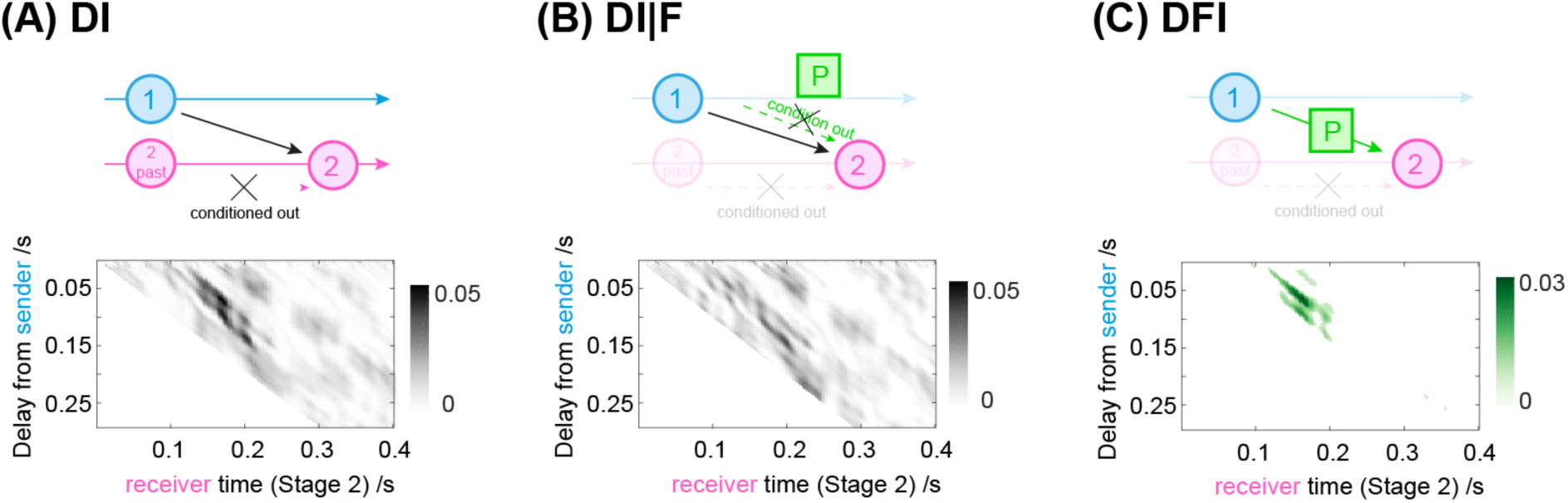
Stage 2: Communications (DFI) of the LSF vs. HSF prediction in the Prediction Network. (A) DI (i.e. transfer entropy) computed between sender source 1 (temporal) and receiver source 2 (frontal network node), for each receiver time point every 2ms between 0 and 400ms post Stage 2 auditory cue onset, and for each sender-receiver communication delay every 2ms between 0 and 300ms. The plot shows DI strength in a typical participant, in the time course of the receiver node (X-axis), as the communication delays from the sender node (Y-axis). DI quantifies all the information communicated from the sender to the receiver nodes, removing information sent from the receiver itself. **(B) DI**|**F** repeats the DI computations, but conditioning out (i.e. removing) the effect of the LSF vs. HSF cue from the DI communications. **(C) DFI** is the difference between DI and DI|F that isolates the specific information communicated between the two network nodes about the LSF vs. HSF predictive cue. For example, temporal node 1 sends the predictive cue P to frontal node 2, with a 50-100 ms delay indicated as a green diagonal in the sender to receiver time x time plot.

**Supplemental Figure S5.**
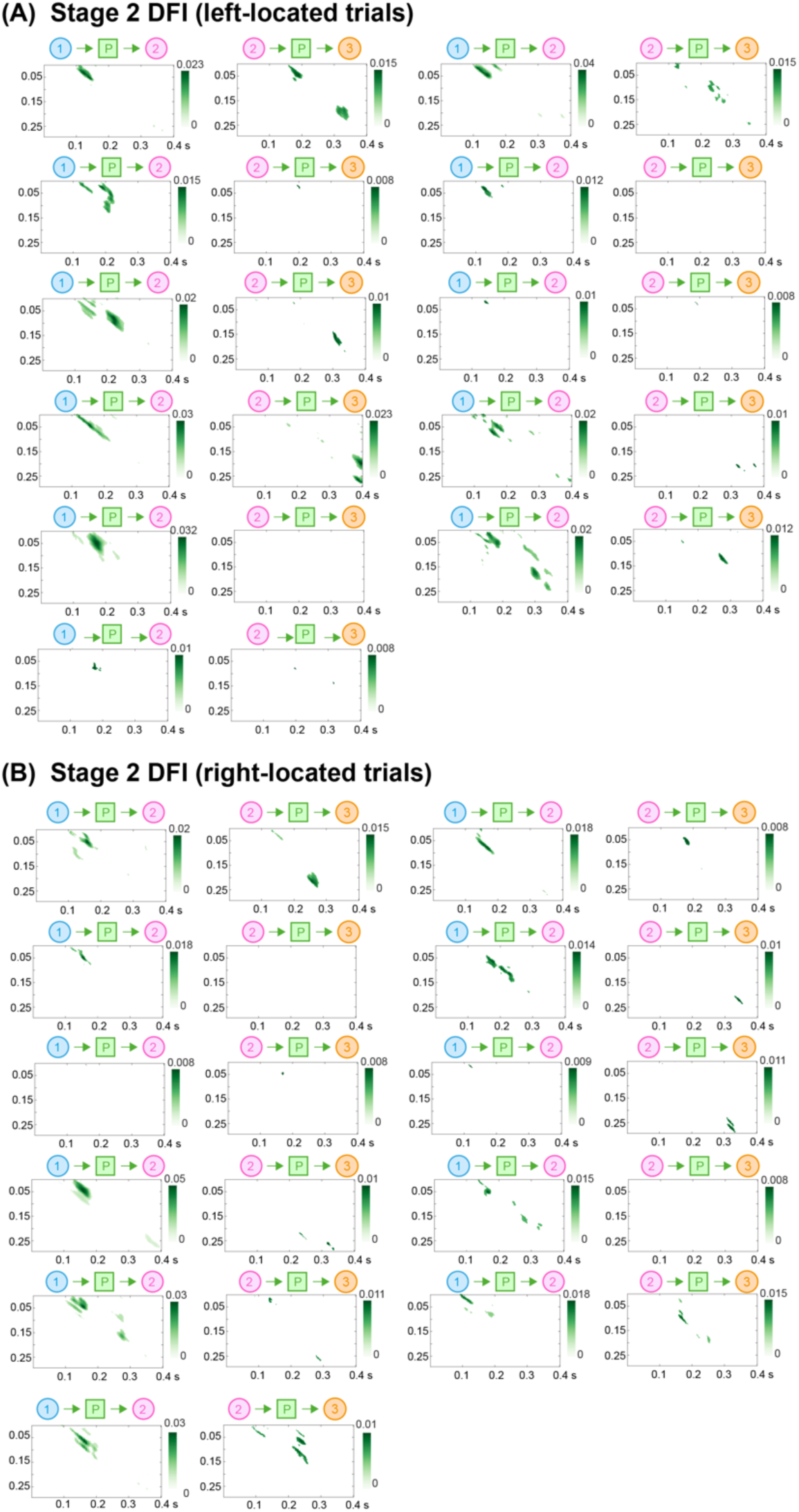
Stage 2: Communications (DFI) of the LSF vs. HSF prediction in the Prediction Network of individual participants. Using DFI, separately for **(A) left-located trials** and **(B) right-located trials**, we computed in each participant the communications of the prediction across network nodes (i.e. 1 (temporal) -> 2 (frontal) and 2 (frontal) -> (3) occipital), every 2 ms between 0 and 400ms post auditory cue onset for the receiver, and every 2 ms communication delay between 0 and 300ms from the sender. These time x time plots represent the significant (FWER-corrected, p<0.05) prediction communications between receiver (X-axis) and sender (Y-axis), where a green diagonal indicates the timing and duration of the prediction communications.

**Supplemental Figure S6.**
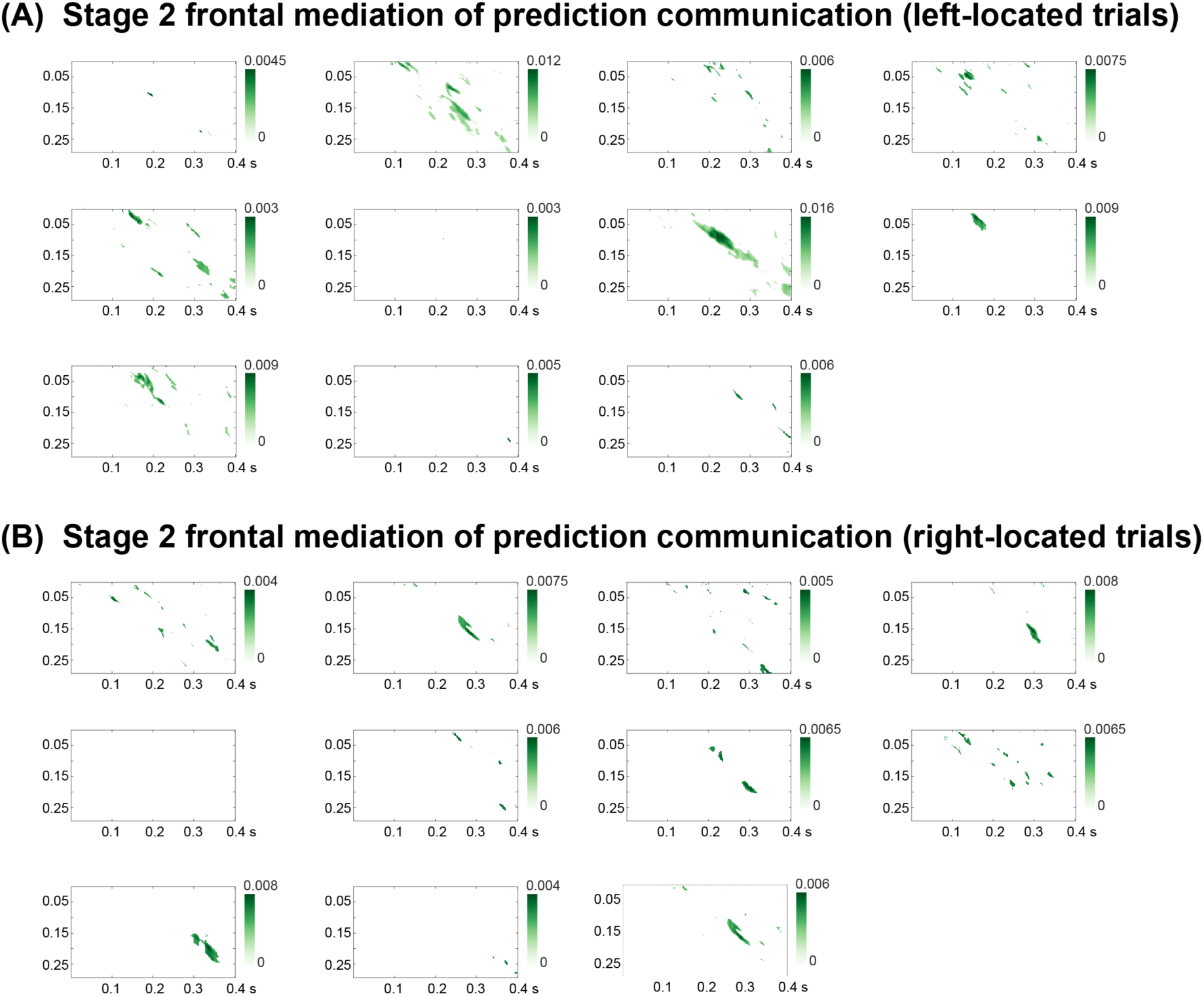
Stage 2: Frontal mediation of prediction communications in the Prediction Network of each participant. Separately for **(A)** left- and **(B)** right-located trials, we computed the difference of temporal to occipital Stage 2 prediction communication, between direct (removing frontal mediation) and mediated (with frontal mediation) DFI (for each receiver time point every 2 ms between 0 and 400ms post auditory cue onset, and for each sender communication delay every 2 ms between 0 and 300ms). Each plot presents the significant (FWER-corrected, p<0.05, one-tailed) frontal mediated Stage communication of the cue between temporal cortex (Y-axis) and occipital cortex (X-axis) network nodes.

**Supplemental Figure S7.**
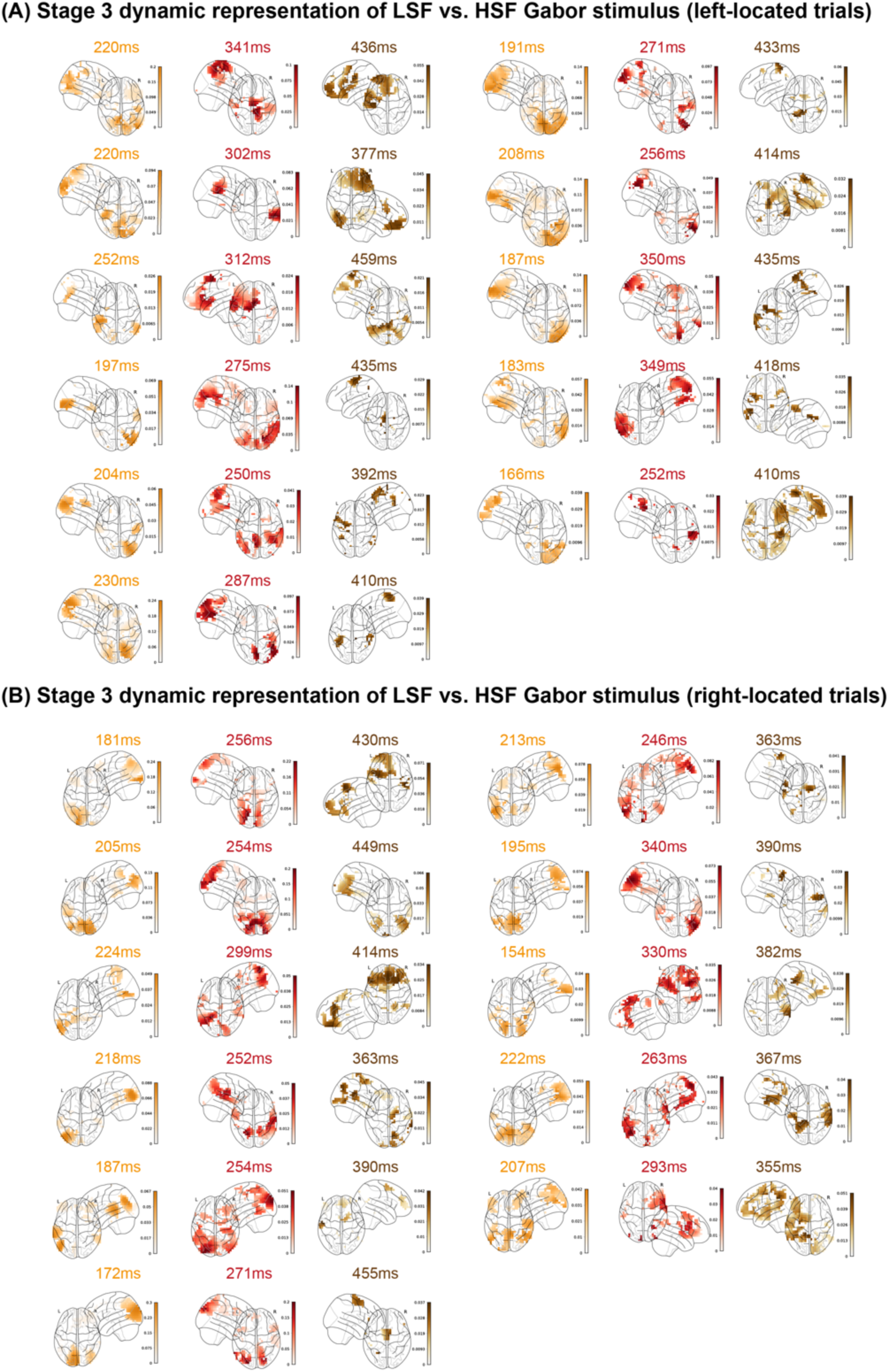
Stage 3: Dynamic representation of the LSF vs. HSF Gabor stimulus in the Categorization Network of each participant. For **(A)** left- and **(B)** right-located trials, we computed MI(<LSF vs. HSF Gabor; Stage 3 MEG_t_>), every 2ms, 0 to 500ms following Gabor onset, on source (see *Methods, Stimulus representations*) and selected the peak MI time of each source. We localized the sources whose MI representation of LSF vs. HSF Gabor with first peak in the 150-250ms time window (start, in orange), the last peak in 250-350ms (midway, in red) and >350ms (end, in brown). Brain plots localize maximal representations of the LSF vs. HSF Gabor in each time window.

**Supplemental Figure S8.**
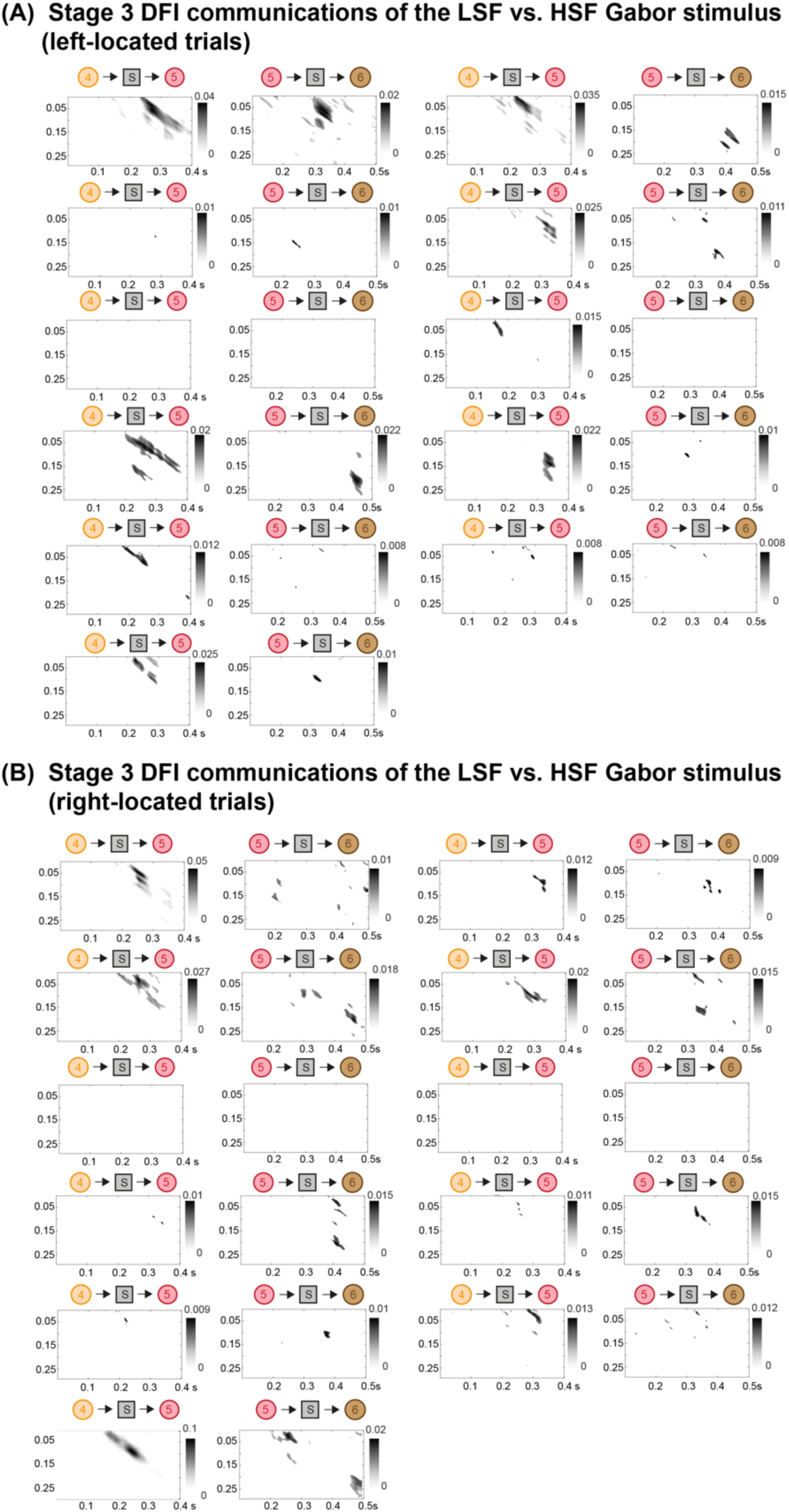
Stage 3: Communications (DFI) of the LSF vs. HSF Gabor stimulus in the Categorization Network of each participant. Separately for **(A)** left- and **(B)** right-located trials, we computed DFI communications between categorization network nodes 4 (occipital) -> 5 (parietal) and 5 (parietal) -> 6 (premotor), every 2ms between 0 and 500ms post Gabor onset for the receiver, and for sender communication delays every 2ms between 0 and 300ms. Each plot presents the significant (FWER-corrected, p<0.05) communications of LSF vs. HSF Gabor stimulus between receiver (X-axis) and sender (Y-axis).

**Supplemental Figure S9.**
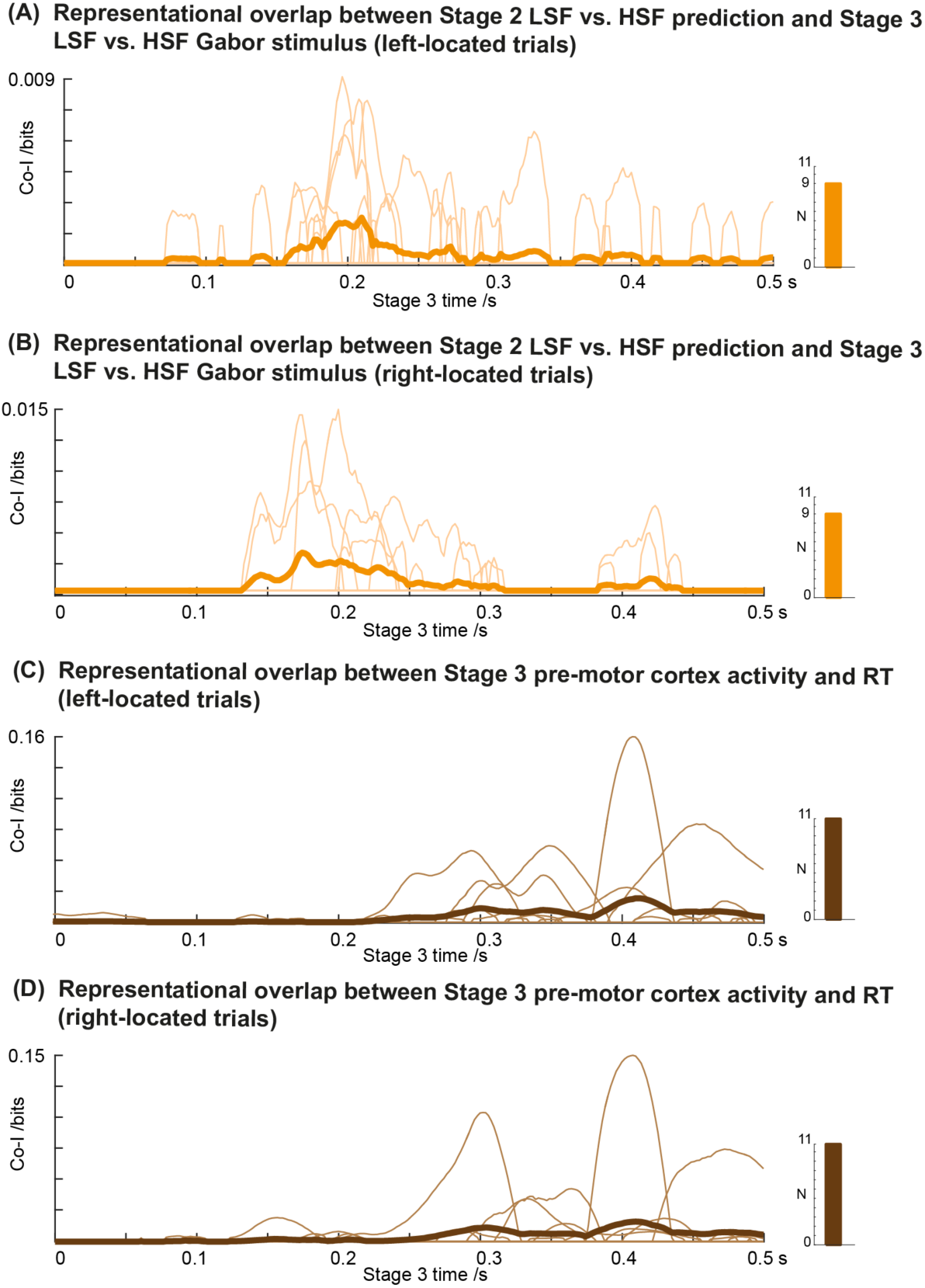
Representational overlaps. Stage 2 to Stage 3: **(A) and (B) Representational overlap between Stage 2 LSF vs. HSF prediction in the Prediction Network and Stage 3 LSF vs. HSF Gabor stimulus in the Categorization Network, left- and right-located trials9.** To measure this representational overlap, first we selected two occipital sources: (1) the source that maximally represents (MI) the Stage 2 LSF vs. HSF prediction; (2) the source that maximally represent (MI) the Stage 3 LSF vs. HSF Gabor stimulus. Then, using these two sources, we computed Co-I(<LSF vs. HSF; Stage 2 Source_1_; Stage 3 Source_2_>), FWER-corrected, *p*<0.05, selecting the maximum Co-I representational overlap across Stage 2 time points. The thick orange curve represents the average prediction-to-categorization representational overlap across participants in occipital cortex, with a peak ∼200ms post-stimulus—independently replicated for left and right-located trials, in 9/11 participants, Bayesian population prevalence = 0.81 [0.53 0.96], MAP [95% HPDI]. The thin orange curves present individual results. **(C) and (D) Representational overlap between Stage 3 pre-motor cortex activity and RT in the Categorization Network (left- and right-located trials)**. To test that prediction modulates both premotor cortex activity and RT, we computed the representational overlap with Co-I(<Predicted vs. Non-predicted trials; Stage 3 premotor MEG; RT>), FWER-corrected, *p*<0.05, using the source with highest MI(<Predicted vs. Non-predicted trials; Stage 3 premotor MEG>). The thick brown curve plots this Co-I averaged across participants, with a peak ∼400ms post Gabor for left- and right-located trials—replicated in all participants for left- and right-located trials, Bayesian population prevalence = 1 [0.77 1] MAP [95% HPDI]. The thin brown curves present individual results.

